# Mapping the Genetic Interaction Network of PARP inhibitor Response

**DOI:** 10.1101/2023.08.19.553986

**Authors:** Danny Simpson, Jia Ling, Yangwode Jing, Britt Adamson

## Abstract

Genetic interactions have long informed our understanding of the coordinated proteins and pathways that respond to DNA damage in mammalian cells, but systematic interrogation of the genetic network underlying that system has yet to be achieved. Towards this goal, we measured 147,153 pairwise interactions among genes implicated in PARP inhibitor (PARPi) response. Evaluating genetic interactions at this scale, with and without exposure to PARPi, revealed hierarchical organization of the pathways and complexes that maintain genome stability during normal growth and defined changes that occur upon accumulation of DNA lesions due to cytotoxic doses of PARPi. We uncovered unexpected relationships among DNA repair genes, including context-specific buffering interactions between the minimally characterized *AUNIP* and BRCA1-A complex genes. Our work thus establishes a foundation for mapping differential genetic interactions in mammalian cells and provides a comprehensive resource for future studies of DNA repair and PARP inhibitors.

## INTRODUCTION

A genetic interaction (GI) quantifies the functional relationship between two genes by measuring the effect of one gene on the phenotype of another. Effects that are less than expected, as determined by individual phenotypes, signify buffering relationships and those that are more than expected represent synergistic effects. Many previous studies have shown that careful examination of GIs can reveal fundamental mechanistic insights, a classic example of which is the GI between *BRCA1* and *53BP1* (*TP53BP1*).^1^ These genes demonstrate a clear buffering relationship, where toxicity associated with *BRCA1* deficiency can be rescued by loss of *53BP1*. Discovery of this GI led to a deep understanding of homologous recombination (HR), where an early nucleolytic processing step is promoted by BRCA1 and suppressed by a host of factors recruited by 53BP1.^2–12^

Genetic interactions can also be exploited for therapeutic purposes. Here, too, *BRCA1* provides an instructive example. Loss-of-function mutations in this gene, as well as other HR genes, predispose individuals to a high incidence of breast and ovarian cancer.^13^ At the cellular level, such perturbations confer sensitivity to loss of poly (ADP-ribose) polymerase (PARP),^14,15^ and observation of these synergistic GIs prompted development of clinical PARP inhibitors (PARPi), now used to treat cancer.^16^ Ostensibly, these drugs kill HR-deficient cancer cells by trapping PARP onto DNA, causing replication stress, and overwhelming the diminished DNA repair capacity of those cells.^17–20^ Further illustrating the importance of GIs in this context, preclinical studies have suggested that alterations in *53BP1* and downstream genes, which buffer the effect of PARPi on HR deficient cells, may underlie clinical resistance.^21^ There is therefore much interest in GIs relevant to PARPi response and how that information may be used to inform new treatment strategies.

Beyond these specific examples, many genetic interactions have informed our understanding of DNA repair and provided guidance for therapeutic development. For this reason alone, a large-scale survey of genetic interactions among human DNA repair genes would have obvious utility, but while previous studies have shown that such efforts are possible,^22–51^ platforms developed for such analysis in human cells^36,43,44,46,48–50^ remain bespoke and have not been extensively applied. Systematic analysis of GIs among human DNA repair genes is therefore lacking. Examining GIs between DNA repair genes also presents a particular challenge, namely that interactions between these genes are predicted to be enriched for context-specific interactions,^51^ which may only be observed after exposure to particular forms of genotoxic stress or in select genetic backgrounds. Global analyses of GIs associated with human DNA repair genes will accordingly require large-scale comparison of such measurements across relevant conditions, demonstration of which has yet to be established. Here, we develop a framework for such work and apply it to map the GI network underlying cellular response to PARPi. Given the chemical-genetic relationship between PARPi and DNA repair, as well as the deep interest in interactions relevant to PARPi-induced genotoxicity, this condition presented the ideal context for the first differential GI map in human cells.

Altogether, we interrogated 147,153 gene pairs under both normal cell growth conditions and in the presence of the PARPi niraparib. Our results reveal a rich topology of functional modules and relationships between DNA repair genes and demonstrate how those relationships change in response to PARPi-induced cell stress. In the context of normal growth, our data provides a high-resolution look at the dense set of relationships among genes responsible for RAD51 filament assembly, stability, and function in HR, including a strong set of novel interactions between genes encoding the molecular motor protein RAD54L and the PSMC3IP-MND1 heterodimer. In the PARPi context, we define interactions with genes encoding the molecular targets of niraparib, *PARP1* and *PARP2*, and identify a striking set of context-specific interactions with a gene only minimally characterized in the context of DNA repair, *AUNIP*. The interaction maps we present here provide a resource for exploring the processes that underlie cellular responses to both PARPi and DNA damage.

## RESULTS

### A dual-sgRNA CRISPRi library targeting pairs of genes involved in PARPi response

We sought to map the network of genes that control genome stability in response to PARP inhibition and evaluate the plasticity of that network in the presence and absence of PARP-induced damage (Figure 1A). To achieve this goal, we deployed CRISPRi, a technique that uses a catalytically inactive SpCas9 fusion protein (dCas9-KRAB) to inhibit transcription at targeted promoters.^52,53^ CRISPRi offered several advantages for this effort. First, among the ∼900 human genes implicated in the DNA damage response,^54^ there are many with strong growth phenotypes. CRISPRi generates partial loss-of-function perturbations and thus limits the severity of interfering with essential genes, allowing robust measurement of phenotypes associated with those genes.^55^ Second, CRISPRi does not alter the sequence of targeted genes and thus avoids toxicity and variability from gene disruption with DNA double-strand breaks.^56,57^ Finally, large-scale mapping of GIs from a single condition across a different set of genes has already been convincingly demonstrated with CRISPRi.^48^

**Figure 1.**
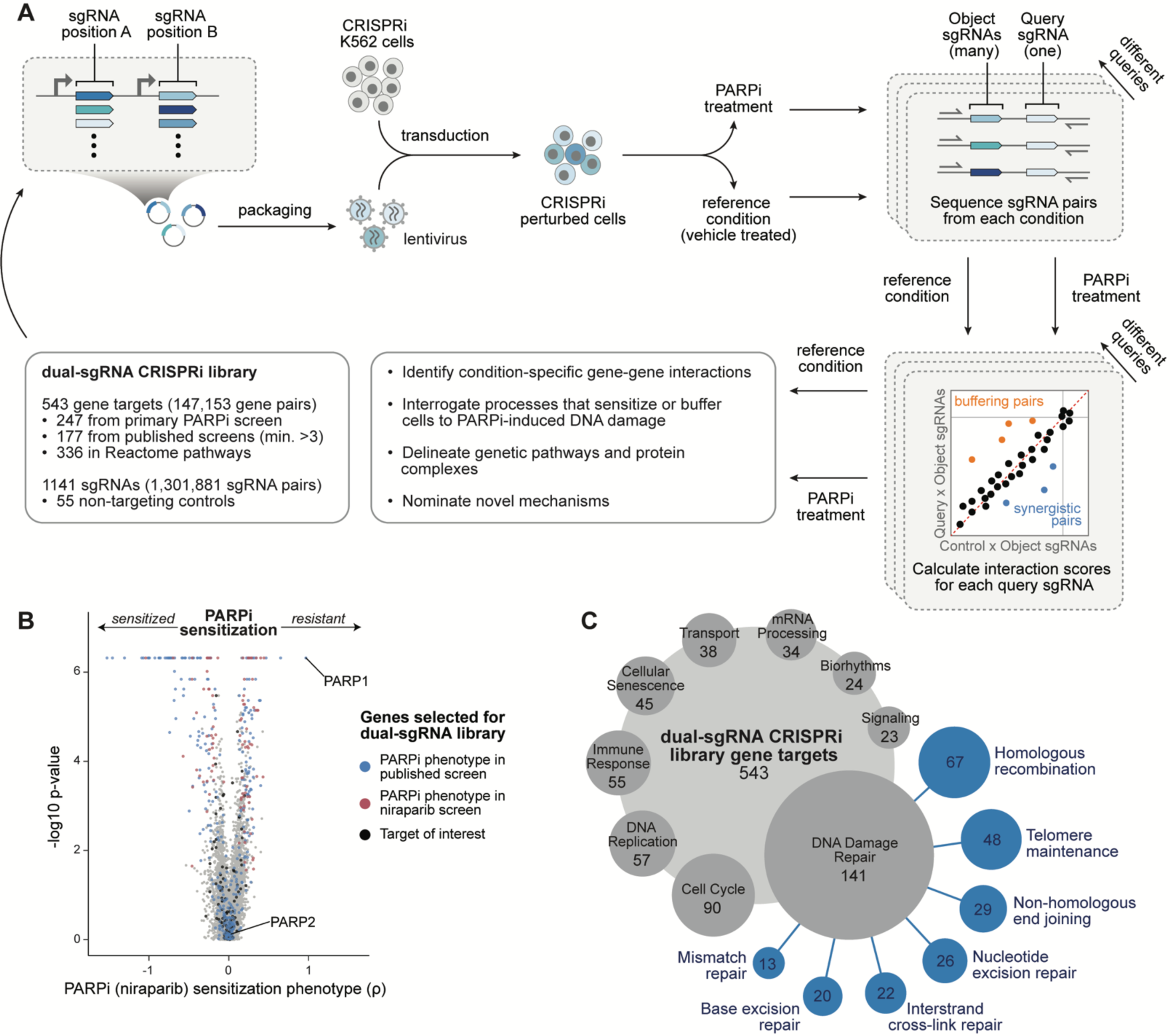
A platform for using CRISPRi to profile context-dependent genetic interactions. (A) Schematic of approach for large-scale genetic interaction mapping with and without exposure to one PARPi, niraparib. (B) Results from genome-scale, single-perturbation CRISPRi screen for genes that cause sensitivity or resistance to niraparib. Genes selected for inclusion in our dual-sgRNA, CRISPRi library are indicated (blue, previously detected in published PARPi screens; red, detected by our genome-scale screen; black, other). P-values determined by MAGeCK (Table S1). (C) Functional annotation of genes included in the dual-sgRNA, CRISPRi library (Table S2; Methods). Genes may appear in multiple categories.

To nominate genes for our map, we conducted a single-perturbation, genome-scale CRISPRi screen using the PARPi niraparib. Results from this screen nominated 247 genes, loss of which caused sensitivity or resistance to niraparib, hereafter referred to as PARPi sensitization (ρ) phenotypes (Figure 1B; Table S1). Among nominated genes were *PARP1*, which conferred the strongest resistance effect, and many genes with known roles in forestalling or repairing PARPi-induced DNA damage (*e.g.*, *BRCA1*, *BARD1*, *DNPH1*, and *PALB2*).^58,59^ To supplement these genes, we also evaluated results from 24 published, single-perturbation CRISPR screens performed with various PARP inhibitors^5,6,8,60–63^, which added another 109 genes. Finally, to ensure comprehensive inclusion of genome stability genes, we selected 187 additional genes from a focused literature review and enriched pathways (Methods), for a total of 543 genes. These genes covered the full spectrum of observed growth phenotypes with and without PARPi exposure, referred to as τ and γ phenotypes respectively (Figure S1A), and demonstrated a high level of functional characterization (*i.e.*, 62% were present in at least one enriched Reactome pathway;^64^ Table S2). Critically, our gene set was also enriched for desired bioprocesses, including DNA repair, DNA replication, and the cell cycle (Figure 1C; Table S2; Methods).

Data from our niraparib screen also allowed us to identify active CRISPRi sgRNAs against our selected genes. From the ∼5 sgRNAs per gene present in the genome-scale library, we chose two per targeted gene (1086 total), specifically selecting those that demonstrated the strongest PARPi sensitization phenotypes with consistent direction, in addition to 55 non-targeting controls. Then, using a two-step cloning strategy with a dual-expression vector, we assembled pairs of these sgRNAs into a “GI library” (Figure S1B; Methods). Notably, because any of our selected sgRNA sequences could appear in either of two positions (A or B) in the expression vector, the final library maximally contained 1,301,881 unique constructs targeting 147,153 gene pairs. Deep sequencing detected all but 502 of these constructs, representing an impressive 99.96% capture rate, with 90% of library elements present within a 6x range and Gini inequality index < 0.28 (Figures S1C and S1D).

### Quantifying genetic interactions in the presence and absence of PARP inhibition

To perform our interaction screen, we transduced our GI library into a K562 CRISPRi cell line,^55^ selected transduced cells, and grew those cells in two replicates with or without exposure to niraparib (Figures 1A and S1E). After nine days, cells were collected and the integrated expression cassettes were amplified from genomic DNA and sequenced to determine the representation of each dual-sgRNA construct at the beginning (T0) and end of the screen (reference and PARPi endpoints) (Figure S1F). Counts from 1,264,760 constructs, representing all those remaining after removing poorly represented elements, were then used to calculate three metrics: reference growth phenotypes (γ); growth phenotypes modified by PARPi exposure (τ); and PARPi sensitization phenotypes (ρ), which directly relate τ to γ (Figure S2A; Methods).^65^ When evaluated individually, as pairs of sgRNAs, or collapsed to gene-level measurements, these phenotypes were highly reproducible, both across replicates and reciprocal orientations in the expression construct (X/Y or Y/X), while non-targeting constructs were, as expected, uncorrelated, with minimal to no evidence of phenotypes (Figures S2B-S2D). Individual gene phenotypes also correlated with data from our primary screen (Figure S2E).

We next used phenotypes from our screen to calculate sgRNA-level interaction scores using an established analytical framework (Methods).^48^ Interaction scores quantify the extent to which phenotypes from pairs of perturbations deviate from expectations set by individual phenotypes. To establish these expectations, we modeled the relationships between individual sgRNA phenotypes and corresponding dual-sgRNA phenotypes for single “query” sgRNAs (Figure 2A). Calculating the difference between measured phenotypes (observed) and model-derived values (expected) then produced GI scores for sgRNA pairs, each of which quantified the direction and strength of the underlying interaction, with positive and negative scores representing buffering and synergistic interactions, respectively. After quality control filtering (Methods), we applied this framework to produce 2,411,208 total sgRNA-level scores, calling those from our reference data γGI scores and those from PARPi-treated cells τGI scores. Among these measurements, we observed evidence of known gene-gene interactions, including buffering interactions between *BRCA1* and *53BP1* as well as synergistic interactions between *BRCA1* and Fanconi anemia (FA) pathway genes (Figure 2A).^66^ To generate gene-level scores, we then averaged all sgRNA-level GI scores targeting the same genes (Table S3).

**Figure 2.**
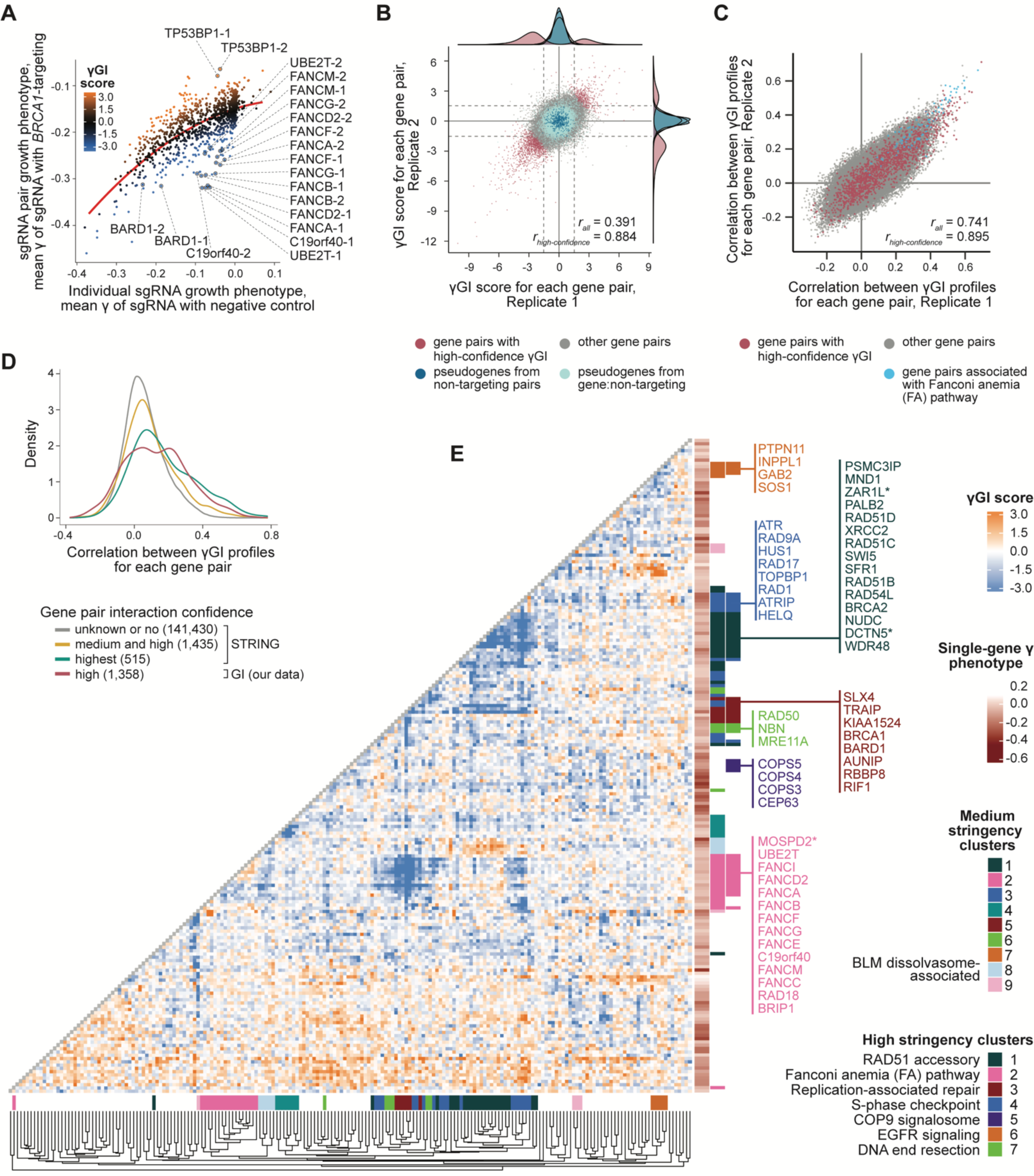
Genetic interactions determined from reference growth (γ) phenotypes reveal a complex interaction landscape among functional DNA repair modules. (A) Example of quantifying sgRNA-level genetic interactions using model-defined expected phenotypes (red) based on the relationship between individual sgRNA phenotypes and the corresponding combinatorial phenotypes with one (*BRCA1*-targeting) query sgRNA. (B) Gene-level γGI score measurements for independent replicates. Dotted lines indicate significance bounds corresponding to FDR ≤ 0.05. Pearson correlations are given for all interactions and for high-confidence interactions. (C) Gene-level γGI profile correlations for independent replicates. Fanconi anemia pathway cluster from E was used to define gene pairs in blue. Correlations as in B. (D) Distribution of γGI profile correlations by confidence. Gray, yellow, and teal indicate confidence in interaction call as determined by STRING evidence score (Methods); red indicates high-confidence interactions identified from GI screen. (E) Reference growth (γGI) interaction map for genes with at least four high-confidence (local FDR ≤ 0.01), supported (≥ 4 contributing scores) interactions. Medium and high stringency clustering (Methods) indicated with color bars. Asterisks indicate genes with a promoter near that of another gene in the same cluster (< 1 kb) and may represent CRISPRi artifacts.

A major advantage of studying genetic interactions in high-throughput is the ability to use model-derived expectations for combinatorial phenotypes. Compared to low-throughput analyses where expected phenotypes are defined only by the two component perturbations, modeling expands the range of observable interaction measurements.^39,48,67^ To clarify this point, consider that our growth-based measurements have an intrinsic limit, complete dropout. As the phenotypes of component perturbations become more severe, a correction must be applied to calculations of expected combinatorial phenotypes to prevent them from dropping below this limit. Plotting phenotypes associated with three sgRNAs for which a simple linear fit properly modeled our reference data illustrates this point (Figure S3A). These data show a relationship between the sgRNA phenotypes of each query guide and the degree of correction (model slope) that must be applied, with guides having more severe phenotypes (*e.g.*, *ZNF574*) requiring more correction. Expanding this analysis showed that calculating expected combinatorial phenotypes from just the component parts (*e.g.*, adding them together) generally requires no correction only when one of those phenotypes is close to zero (Figure S3B). Indeed, applying this simpler approach to our data and averaging all resulting genetic interaction scores for each gene showed stronger expected combinatorial phenotypes than with a model framework, resulting in a bias towards calling buffering interactions which increased as gene growth phenotypes became more severe (Figure S3C). Use of a modeling framework corrects for this effect and, consistent with previous observations,^48^ we found that a quadratic model outperformed a linear model for 86% of query sgRNAs (F-test), further highlighting the advantage of high-throughput analysis.

Altogether, we recovered 143,354 gene-level GI scores from our reference condition, representing 139,259 novel GI measurements compared to previous analysis,^48^ and 145,500 from PARPi-treated cells (>97% of targeted gene pairs in each case; Table S3). Our data thus dramatically increases existing knowledge of genetic interactions relevant to both conditions. Providing confidence in these measurements, we validated several in low-throughput and found a strong correlation with screen values (Figure S3D). Additionally, to guide exploration of these data, we examined and annotated technical features associated with each measurement (Figures S3E-S3I; Table S3; Methods). Finally, using permutation testing, we determined the null distributions of these GI scores and identified 1358 and 1676 high-confidence buffering and synergistic interactions (local FDR ≤ 0.01) in the reference and PARPi conditions, respectively, as well as 10,139 and 11,976 significant interactions (FDR ≤ 0.05; Table S3; Methods). We call these significant interactions γGIs and τGIs (distinguishing from γGI and τGI *scores*, which refer to measurements regardless of significance).

### High-resolution mapping of genome stability genes and related processes

Our GI library measures >1.3 million sgRNA-level interactions per condition but because genetic interactions with a buffering or synergistic effect are rare,^68^ the vast majority of our measurements were not expected to show such relationships. Replicate sgRNA- and gene-level interaction scores were thus only mildly correlated when evaluated altogether (Figures 2B, S4A, and S4B). Nevertheless, confirming that our data contained reproducible signals, replicate correlations sharply increased when restricted to only significant or high-confidence interactions, which appropriately represented a minority of measured scores (0.9% high-confidence γGIs, 7.1% significant γGIs; 1.1% high-confidence τGIs, 8.2% significant τGIs). Genetic interactions are also not evenly distributed across genes; indeed, across established networks, a small number of genes have been found to have an unusually high number of interactions.^33^ These genes often encode conserved, multifunctional, highly expressed and abundant proteins.^39^ A handful of genes in our dataset demonstrated many more γGIs and τGIs than average; specifically, the 5% most highly interacting genes were involved in 33% of all significant reference interactions (3,341) and 34% of all significant PARPi interactions (4,084). Weak correlation between the number of significant interactions per gene and growth phenotypes also suggested that highly interacting genes may be more likely than lowly interacting genes to be essential for growth in the corresponding condition (Figures S4C and S4D).

Another advantage of measuring genetic interactions in high-throughput is the ability to explore genetic interaction profiles; *i.e.*, arrays of interaction measurements made for one gene across many others. These profiles serve as quantitative representations of gene function and, when evaluated at scale, can identify groups of genes with broadly similar functions and determine how those groups, or gene “modules”, interact. Correlations between gene-level γGI profiles from our data were highly reproducible, particularly among gene pairs with high-confidence interactions, and tended to be higher between gene pairs encoding physically interacting proteins (Figures 2C and 2D; Methods). Notably, while not independent from GI scores, these profiles do provide complementary information. For example, while only a minority of gene pairs associated with the FA pathway demonstrated significant γGI scores (22 of 91), many FA interaction profiles were among the most highly correlated of any pairs in our library (Figure 2C). Our data can thus be used in multiple ways to quantify functional and physical relationships, and highlighting the potential for discovery, clustering interaction profiles from our reference data revealed many groups of functionally related genes (Table S4).

To aid exploration of gene clusters within our data, we subset our reference data to include only genes (200) with an excess number of supported, high-confidence gene-level γGI scores, which we defined as ≥ 4 such scores, the median number among genes with at least one (Figure 2E; Methods). This subsetting captured 73% of high-confidence γGIs (996) and maintained sufficient information to obtain clustering similar to the full data set while also reducing information density for visualization (Figure S4E). High stringency clustering identified seven clusters, including ones composed of RAD51 accessory genes, FA pathway genes, S-phase checkpoint genes, DNA end resection genes, and one with genes loosely connected through roles in repair after replication stress. Reducing stringency expanded the number of genes within these modules and added three groups, including one containing genes associated with the BLM dissolvasome complex. Altogether, clusters revealed interrelatedness among and within gene modules. Genes within the FA pathway cluster, for example, were largely synergistic with those in the replication-associated repair cluster, while structure within the FA cluster highlighted functional differentiation among member genes, including separating out *FAAP24 (C19orf40)* and *FANCM*, reflecting the modularity of FA subcomplexes.^69–71^ Notably, we also observed interactions between groups of genes not primarily associated with DNA repair, for example, buffering relationships between EGFR and Ras-MAPK signaling genes,^72^ and interactions between individual genes and modules, such as a novel synergistic relationship between COP9 signalosome genes and the kinetochore-associated *CENPF* (Figure 2E; Tables S3 and S4).^73^ Clustering applied to our complete reference map identified 24 total modules (Table S4), forming a complex network of interactions across a range of biological processes, but with specific focus on DNA repair mechanisms relevant to normal cell growth.

### Quantifying a dense set of functional relationships among homologous recombination genes

One of the gene modules identified within our subsetted reference map, the RAD51 accessory cluster, was composed primarily of genes with accessory functions in homologous recombination (Figure 2E). HR is a complex repair process that uses template-dependent DNA synthesis to repair broken DNA and rescue stalled replication forks.^74^ The RAD51 recombinase, which polymerizes on single-stranded DNA exposed at DNA lesions, promotes the search for and invasion of an intact homologous template. This protein is therefore essential for homologous recombination but, indicative of the complexity of HR, does not act alone. Rather, RAD51 is aided by various accessory proteins. High stringency clustering of our subsetted γGI profiles isolated genes encoding such proteins with remarkable precision. Within a single module, we observed genes whose encoded products (a) recruit RAD51 to breaks (*BRCA2*, *PALB2*);^75^ (b) facilitate and stabilize RAD51 nucleoprotein assembly (*RAD51B*, *RAD51C*, *RAD51D*, *XRCC2*, *SWI5*, *SFR1*, *RAD54L*);^76–79^ (c) stabilize filaments of RAD51 and its paralog *DMC1* in meiosis (*PSMC3IP*, *MND1*);^80,81^ (d) promote formation of the three-stranded displacement loop (D-loop) formed after invasion of the homologous DNA template (*PSMC3IP*, *MND1, RAD54L, WDR48*);^82,83^ and (e) facilitate RAD51 removal after D-loop formation (*RAD54L*).^84^

Demonstrating the relatedness of genes within this cluster, many were observed to encode subunits of different protein complexes, including the SWI5-SFR1,^77,85^ PSMC3IP-MND1,^86^ and BCDX2 (RAD51B, RAD51C, RAD51D, XRCC2) complexes.^87^ Examination of experimentally validated physical interactions (from STRING)^88^ among cluster genes, however, revealed far less relatedness then our γGIs, which connected 72% of gene pairs with a significant, negative γGI (56 of 78), overall suggesting wide-spread, functional redundancy between RAD51 accessory proteins and complexes (Figures 3A and 3B). Among these interactions were γGIs between *RAD54L* and *PSMC3IP-MND1* (Figure 3C), two of the ten strongest synergistic relationships in the entire dataset (Table S3). These interactions were of particular interest as *PSMC3IP* and *MND1* have only recently been implicated in somatic recombination.^89–92^ Validation experiments in two cell lines (K562 and diploid, non-transformed RPE1 cells) confirmed that, while individual loss of *PSMC3IP*, *MND1*, or *RAD54L* minimally affects cell growth, combined loss of *RAD54L* with either *PSMC3IP* or *MND1* causes strong growth defects (Figures 3D, 3E, and S4F), supporting a role for PSMC3IP-MND1 in somatic recombination and demonstrating a strong relationship with RAD54L, a DNA-dependent ATPase that regulates RAD51 at the different stages of filament stabilization, D-loop formation, and removal from paired strands.^93^

**Figure 3.**
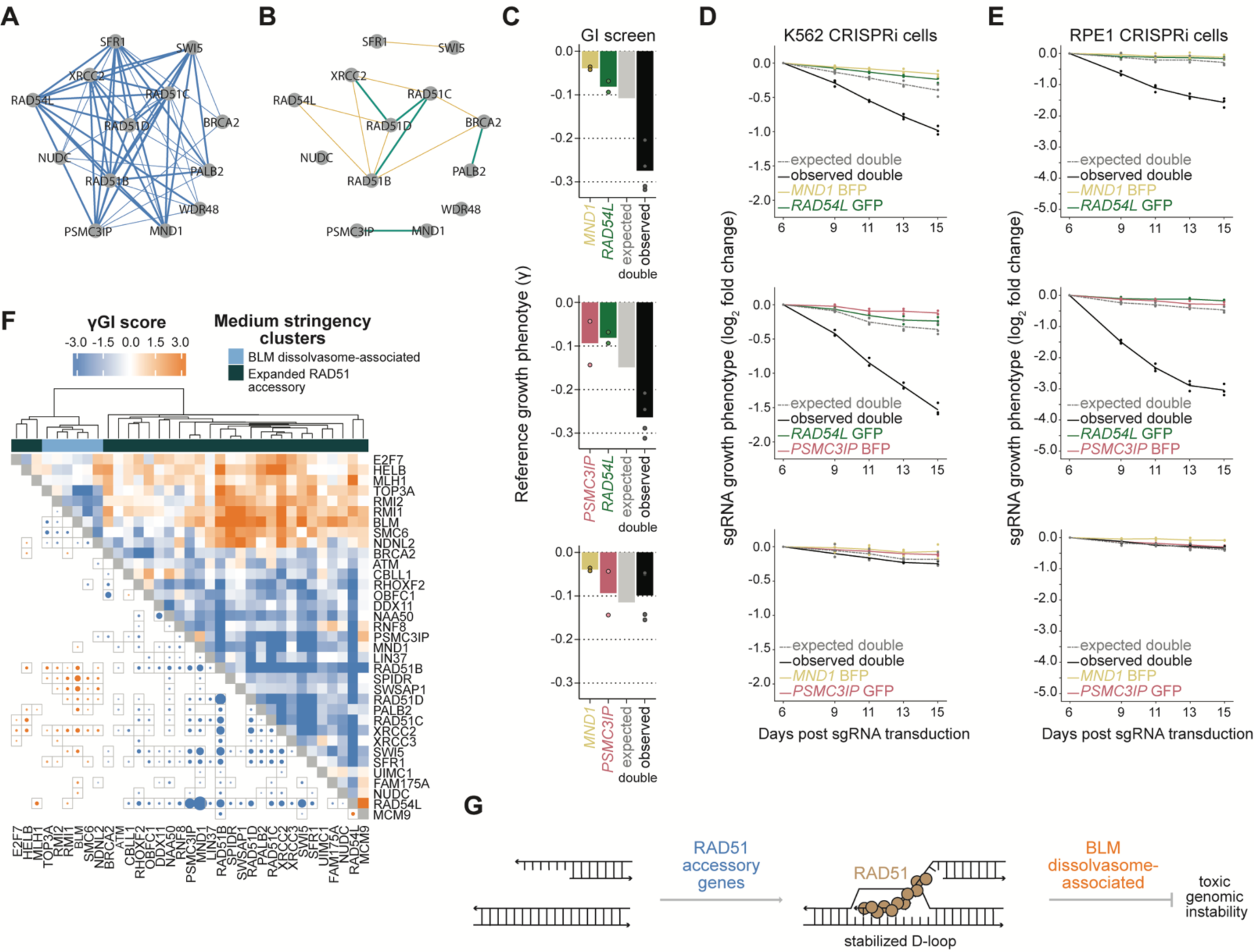
Analysis of a dense set of genetic interactions among RAD51 accessory genes. (A) Network of genetic interactions among RAD51 accessory genes. Edges indicate significant (thin) and high-confidence (thick) interactions; all interactions are synergistic. (B) Network of physical interactions among RAD51 accessory genes. Edges indicate medium-(yellow), high-(yellow), and highest-confidence (teal) interactions defined by STRING (Methods). (C) Observed reference growth (γ) phenotypes for *MND1* (yellow), *PSMC3IP* (red), *RAD54L* (green), as well as observed (black) and model-defined, expected combinatorial phenotypes (gray). sgRNA-level phenotypes (individual or combinatorial) contributing to each gene-level phenotype are shown as dots. (D-E) Growth-based validation experiments in K562 CRISPRi (D) and RPE1 CRISPRi (E) cells for *MND1*, *PSMC3IP*, and *RAD54L* interactions (Methods). Experiments were performed in triplicate, and in each replicate, expected combinatorial phenotype (double) were defined as the sum of the individual contributing phenotypes. Lines indicate replicate averages. Colors as in C. (F) Genetic interactions from reference map (γGI) among genes within an expanded RAD51 accessory module and the BLM dissolvasome complex. Included genes were identified by a medium stringency clustering of all genes in the GI library (Methods; Table S4). The lower left triangle of the heatmap shows only significant interactions, with the strength of significance indicated by size. (G) Roles of RAD51 accessory factors and BLM dissolvasome complex.

We next examined our expanded cluster of RAD51 accessory genes from less stringent clustering of our full data set (Figure 3F; Table S4; Methods). Among genes added to the module from this analysis were two associated with the human Shu complex, *SWSAP1* and *SPIDR*. Similar to other RAD51 accessory factors, the Shu complex promotes RAD51 filament assembly, but unlike others, these factors support recombination between homologous chromosomes and not intra-chromosomally.^94,95^ Indicative of a difference in function, we observed few significant interactions between *SWSAP1* or *SPIDR* and other RAD51 accessory genes; however, examining full γGI profiles revealed buffering interactions with BLM dissolvasome associated genes, similar to other RAD51 accessory cluster genes (Figures 3F, S4G, and S4H).^96,97^ The BLM dissolvasome processes recombination intermediates, and loss of the complex leads to increased sister chromatid exchanges, interhomolog crossing over, and cellular toxicity.^98^ Physical and functional interactions between the BLM helicase and specific RAD51 accessory factors (RAD51D, XRCC2, XRCC3, SWS1, SWSAP1, SPIDR, RAD54L) have been reported previously,^95,99–103^ with buffering interactions attributed to a combination of promoting and forestalling the creation of toxic recombination intermediates (Figure 3G). Our data highlights such antagonism between cluster genes. Moreover, novel relationships with SMC5-SMC6 complex components *SMC6* and *NSMCE3* (*NDNL2*)^104^ support a role for this complex in limiting toxic recombination intermediates alongside the BLM dissolvasome in human cells. Altogether, the known and unknown relationships revealed by these results demonstrate the depth of functional information provided by our data.

### Differential genetic interactions reveal a complex landscape of PARPi-induced damage

DNA repair mechanisms respond to DNA damage in a lesion-specific way,^54^ with dedicated sensors and effectors activated by various chemical and structural irregularities. Assuming that lesions arising during normal cell growth differ from those induced by genotoxic stress in either form or frequency, then genetic interactions among DNA repair genes should also be different. To quantify interactions that differ in the presence and absence of niraparib, we initially turned to PARPi sensitization (ρ) phenotypes and applied a model-based framework as described above for γ and τ phenotypes. Compared to γ phenotypes, however, these models generally had poorer explanatory power, which we attribute primarily to the increased variability from using two endpoint measurements (Figures S2B and S2C), though sgRNAs with strong individual ρ phenotypes were particularly susceptible to poor model performance and also contributed to overall model degradation (Methods). Moving away from using ρ phenotypes and taking inspiration from approaches applied in yeast,^34,38,51^ we defined differential interaction scores, termed νGI scores, for each sgRNA pair by subtracting its γGI scores from the corresponding τGI scores. We then averaged those values to produce νGI scores for each gene pair (Table S3; Methods). Compared to τGI scores called from PARPi-treated cells alone, νGI scores isolate PARPi-specific effects from those of gene perturbation under reference conditions in a way analogous to the underlying phenotypes, where ρ phenotypes separate effects due to PARPi sensitization (Figure S2A).

As with γGI and τGI measurements, we used permutation testing to assign significance to νGI scores, defining “hit” νGIs, or simply νGIs, as those with FDR ≤ 0.2 to account for increased noise from comparing across conditions (Methods). These calculations identified 960 νGIs, 439 positive and 521 negative (Figure 4A; Table S3). Examining these differential interactions, we found that the number of νGIs per gene moderately correlated with the magnitude of sensitivity or resistance to PARPi (Figure S4I), indicating that genes with many νGIs are more likely to be central to PARPi response. Affirming this observation, genes implicated in preventing the formation of PARPi sensitizing DNA lesions (*e.g.*, *DNPH1*, *POLB*, *LIG3*)^59,105–107,20^ or promoting repair of the toxic consequences of PARP trapping (*e.g.*, *FANCA*, *RBBP8*, *RAD18*)^108–110^ were among the most highly interacting genes in the νGI space.

**Figure 4.**
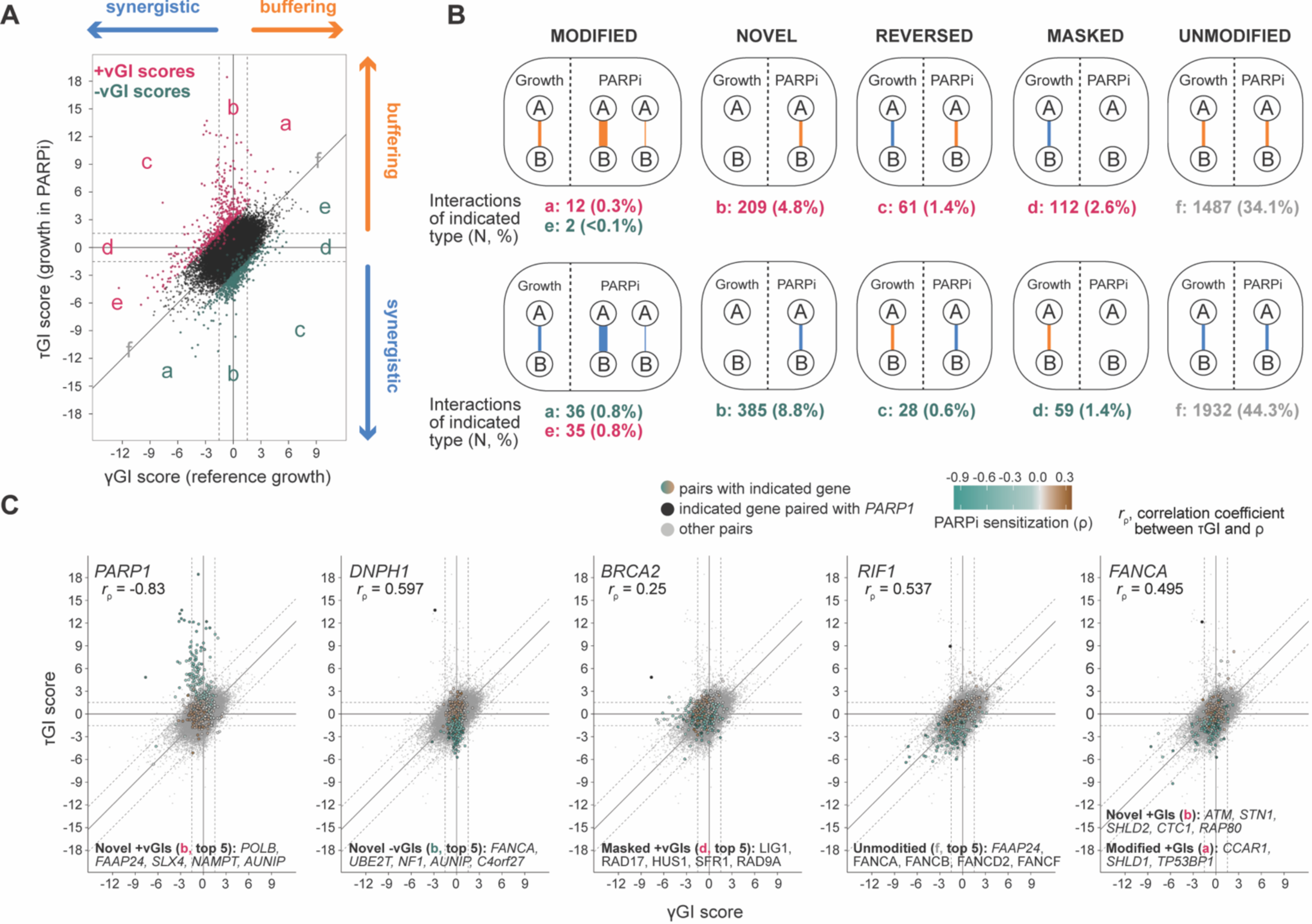
Differential interaction scores reveal context-dependent gene-gene interactions. (A) Topology of γGI and τGI scores with differential interactions indicated (νGI, pink). Dotted lines indicate significance bounds corresponding to FDR ≤ 0.05. Letters indicate regions corresponding to different interaction categories defined in B. (B) Schematics of νGI categories. A small number of νGI hits were uncharacterized (21), with no significant γGI or τGI (Methods). (C) Topologies of γGI and τGI scores for individual genes. Points associated with the indicated gene are colored by the PARPi sensitization phenotype (ρ) of the corresponding partner gene; points not associated with the indicated gene are gray; points representing scores of the indicated genes paired with *PARP1* are black (excepting the *PARP1* panel). Dotted lines indicate significance bounds corresponding to FDR ≤ 0.05 for γGI and τGI, with the diagonal set indicating bounds defining νGI hits at FDR ≤ 0.20. Pearson correlations are shown between individual ρ phenotypes and τ interaction scores.

Next, guided by information from all measurements, we classified νGI hits into four types: “novel”, “masked”, “reversed”, or “modified” (Figure 4B).^51^ Novel interactions were those with a significant τGI score but no significant γGI score. These interactions quantify relationships observed only in the presence of PARPi. Masked interactions were the opposite of novel, meaning that they quantified relationships disrupted by the presence of PARPi. Reversed interactions were those with significant γGI and τGI scores of opposite signs such that the nature of those relationships (*i.e.*, buffering or synergistic) flipped across conditions, and modified interactions were those with significant γGI and τGI scores of the same sign but modified magnitude. In addition to these classifications, many gene pairs demonstrated significant γGI and τGI scores that were unchanged across conditions according to our thresholds. We interpret these “unmodified” interactions as those not affected by the presence of PARPi. Finally, to aid exploration of interactions that were not νGI hits but for which either a significant γGI or τGI score was observed, we added categories of “possibly novel”, “possibly masked”, and “possibly reversed” (Figure S5A; Tables S3 and S5).

For several genes, examining νGI scores across all partner genes allowed inference of gene function in relation to PARP inhibition (Figure 4C). A clear example was *PARP1*, a molecular target of niraparib. Loss of *PARP1* prevents PARPi-induced trapping of the encoded enzyme on DNA and thus forestalls cellular toxicity. Any perturbation conferring sensitivity to PARPi through PARP1 trapping should therefore have a buffering interaction with *PARP1* in the presence of the drug but not necessarily in the absence. Suggesting that such perturbations were common in our library, an overwhelming number of νGIs associated with *PARP1* were positive and novel (92 of 135). Moreover, we found that τGI scores associated with *PARP1* were inversely correlated with the PARPi sensitization (ρ) phenotypes of partner genes. A second example was *DNPH1*, which, in contrast to *PARP1*, had mainly novel and negative νGIs (30 of 36; Figure 4C). The interaction topology of *DNPH1* showed little connection to other library genes during normal growth but many synergistic relationships in the presence of PARPi, consistent with the characterized function of *DNPH1* in forestalling PARPi-trapping by preventing incorporation of aberrant nucleotides into the genome. These observations further suggest that such nucleotides, on their own, do not elicit replication stress.^59^ Notably, τGI scores associated with *DNPH1* also correlated with the PARPi sensitization of partner genes, which, as illustrated by these two examples, was often true for novel interaction enriched genes (Table S5). In other examples of νGI topologies, we observed ones enriched for other types of interactions, such as *BRCA2* with masked interactions across S-phase checkpoint genes (*i.e., RAD17*, *HUS1*, *RAD9A*, and *RAD1*), *RIF1* with unmodified interactions with FA pathway genes, and more complex topologies indicative of a diversity of functional relationships, such as *FANCA*. Examining these topologies also revealed specific interactions of particular interest, several of which we explore in greater depth below.

### Combined loss of RNase H2 and *PARP2* synergistically kills cells with and without PARPi

While many *PARP1* associated νGIs could be attributed to loss of PARP1 trapping, as described above (Figure 4C), we observed relatively few interactions for *PARP2*, which is responsible for ∼10% of cellular break-induced PARylation relative to the ∼90% from *PARP1* (Figure S5B). ^111^ Because the catalytic domains of PARP1 and PARP2 share a high degree of homology, most PARP inhibitors, including niraparib, have a similar potency against both enzymes.^17,112^ Niraparib has also been reported to retain both PARP1 and PARP2 on DNA, with the effect on PARP2 described as PARP1-independent.^113^ Mechanistic inferences from the apparent lack of *PARP2* associated GIs would require careful consideration of the relationship between gene dosage, PARylation, and enzyme trapping, but examination of the νGI topology nevertheless revealed interactions of relevance to both enzymes, specifically unmodified GIs between *PARP2* and three genes previously connected to *PARP1*, namely *RNASEH2A*, *RNASEH2B*, and *RNASEH2C* (Figures 5A-5C; Table S3).^61^

**Figure 5.**
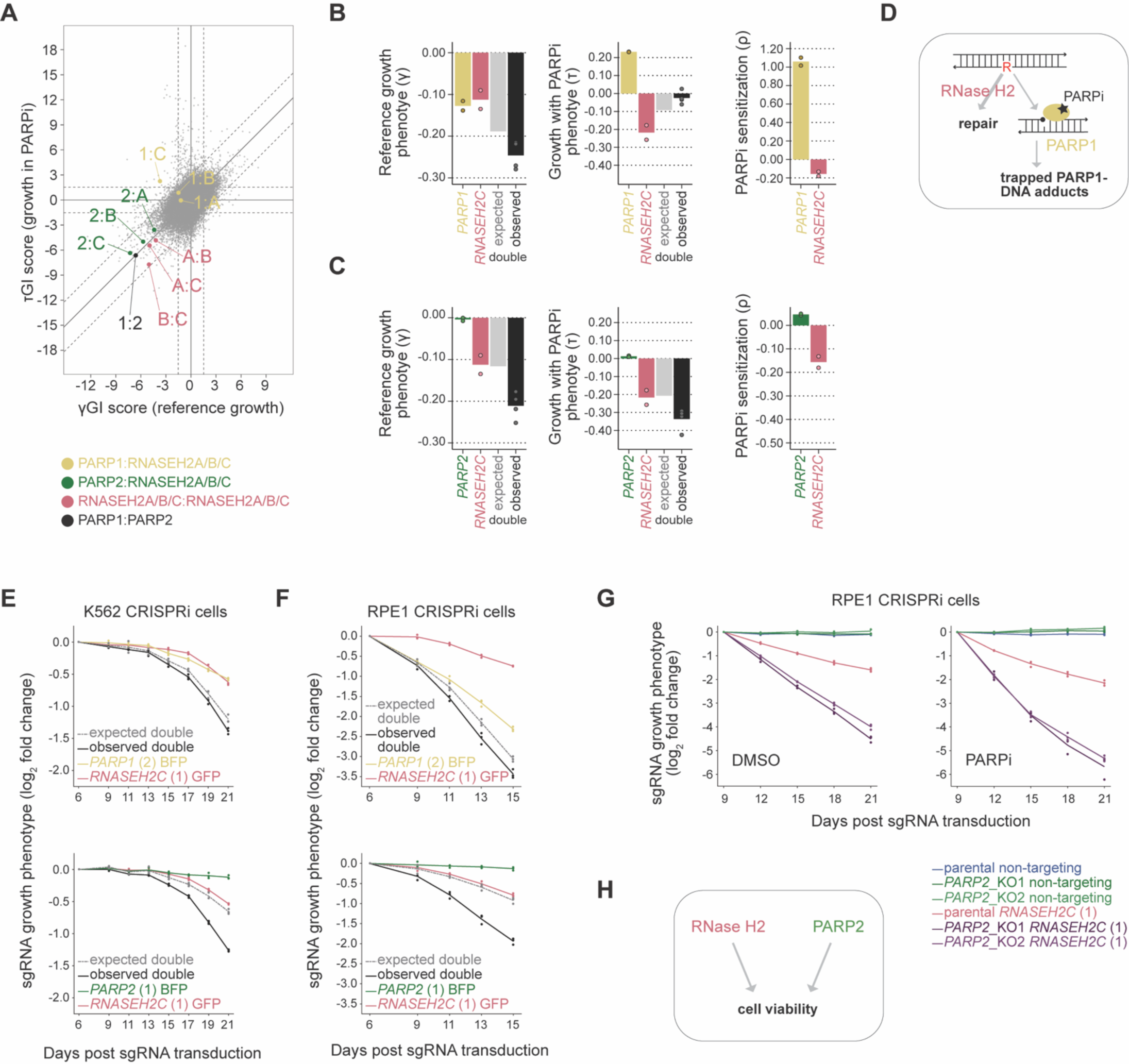
Combined loss of RNase H2 and *PARP2* synergistically kills cells with and without PARPi. (A) Topology of γGI and τGI scores highlighting interactions between *RNASEH2A*, *RNASEH2B*, and *RNASEH2C* with *PARP1* (yellow), *PARP2* (green), or themselves (red). Dotted lines as in Figure 4C. (B-C) Phenotypes (γ, τ, ρ) associated with single or combinatorial perturbation of *RNASEH2C* and *PARP1* (B) or *RNASEH2C* and *PARP2* (C). Model-defined, expected combinatorial phenotypes are shown for γ and τ. (D) Previously established model for the chemical-genetic interactions between RNase H2 and PARPi.^61^ (E-F) Validation of *PARP1*:*RNASEH2C* (top) and *PARP2*:*RNASEH2C* (bottom) interactions in K562 CRISPRi (E) and RPE1 CRISPRi (F) cell lines. As in Figure 3D. (G) Validation of *PARP2*:*RNASHE2C* interaction in two *PARP2* knockout RPE1 CRISPRi clones treated with vehicle (DMSO, top) or PARPi (bottom). As in Figure 3D. (H) *PARP2* and Rnase H2 are synthetic lethal due to parallel, redundant function.

*RNASEH2A*, *RNASEH2B*, and *RNASEH2C* encode three subunits of RNase H2, an enzymatic complex responsible for removing misincorporated ribonucleotides and RNA/DNA hybrids from the genome.^114^ A synthetic lethal chemical-genetic interaction was previously demonstrated between loss of this complex and PARPi.^61^ This interaction was attributed to toxicity from trapping PARP1 on DNA lesions generated in the absence of RNase H2 at sites of ribonucleotide misincorporation (Figure 5D). Consistent with this model, we found that loss of RNase H2 sensitized cells to niraparib (Figure 5B). We also observed a negative, synergistic relationship between *RNASEH2C* and *PARP1* during normal growth, as would be expected if PARP1 contributes to repair of RNase H2-suppressed lesions. Further, reflecting the switch to a toxic relationship between the presence of PARP1 and loss of RNase H2 in the context of PARP inhibition, the interaction between these genes reversed with niraparib (Figures 5A and 5B). By contrast, interactions between *PARP2* and RNase H2 genes were unmodified by niraparib, such that the relationship between these genes appeared unchanged by the inhibition and/or trapping of either enzyme (Figures 5A and 5C).

Focused experiments in two different cell lines (K562 and RPE1) validated the γGI results from our screen, specifically confirming synergism between *RNASEH2C* and *PARP2*, as well as a mild interaction of *RNASEH2C* with *PARP1* (Figures 5E, 5F, S5C, and S5D). Notably, efficient repression by *PARP1*-targeting sgRNAs showed that the weaker interaction observed for *PARP1* was not due to insufficient knockdown (Figure S5E). However, as discussed above (Figure S3C), since PARP1 and RNASEH2C individually demonstrated cell growth phenotypes, the additive approach used here potentially overestimated the expected combinatorial phenotypes and thus lessened the calculated interaction strength (Methods). We next reasoned that, due to partial knockdown from CRISPRi, the synergistic increase in cellular toxicity from loss of *PARP2* and RNase H2 could represent either parallel loss of redundant functions or exacerbated loss of the same activity. Consistent with this logic, genes encoding subunits of RNase H2 demonstrated unmodified interactions with each other, probably due to synergistic destabilization of the complex (Figure 5A).^114–116^ To distinguish between pathway architectures, we generated two *PARP2*-knockout clones in RPE1 cells and evaluated growth with and without perturbation of *RNASEH2C* (Figures 5G and S5F). Here again, we observed a synergistic increase in cellular toxicity, showing the related functions of *PARP2* and RNase H2 either work in parallel or do not completely overlap (Figure 5H). These data showcase the complex roles of *PARP* genes in supporting cell growth and implicate *PARP2* in an important, but as yet uncharacterized, cellular process with RNase H2.

### Differential genetic interactions extensively connect *AUNIP* to the PARPi response

While *PARP1* had by far the largest number of differential interactions in our dataset (135 νGIs), a second gene, *AUNIP*, stood out both in terms of number of νGIs (57, 8 positive, 49 negative) and strength of those measurements; indeed, among the 26 gene pairs with the strongest negative νGIs (top 5%), *AUNIP* was involved in 12 (46%) (Figures 6A and 6B). We interpret the magnitude of νGIs to reflect the extent to which individual functional relationships differ between conditions. As such, observation of many strong νGIs, backed by a relative dearth of unmodified interactions (7), suggests that in relation to other genes, the role of *AUNIP* changed dramatically upon PARP inhibition. Even compared to genes with highly similar PARPi sensitization phenotypes, *AUNIP* had a larger number of strong νGI (Figure 6A). Thus, *AUNIP* stands out among PARPi-sensitizing genes as being particularly context-specific. Similar to *PARP1* and *DNPH1* νGIs topologies (Figure 4C), many *AUNIP*-associated νGIs were novel (43, 75%) and τGI scores were correlated with the single-gene ρ phenotypes of partner genes (Figure 6B); however, different than the inferred function of *PARP1* and *DNPH1*, we reason that the many novel νGIs observed for *AUNIP* are likely explained by a role in context-specific repair, as *AUNIP* demonstrated high-confidence γGIs with several FA genes (7).

**Figure 6.**
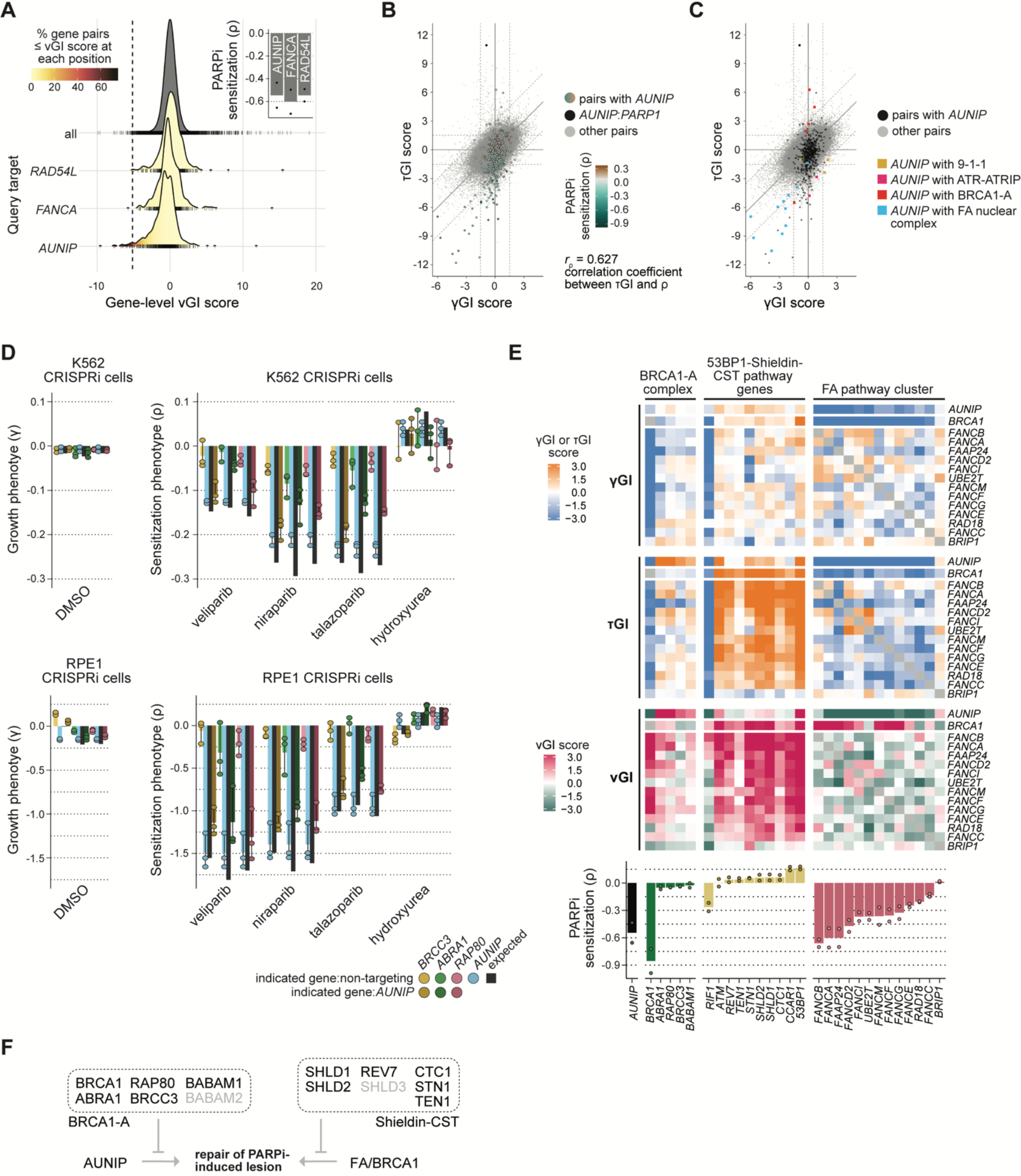
AUNIP has a PARPi-specific buffering interaction with the BRCA1-A complex. (A) Distribution of νGI scores for all gene pairs (top) and for pairs involving *RAD54L*, *FANCA*, or *AUNIP*. Dotted line corresponds to the top 5% of strongest negative νGI hits; rug plots show individual interactions in each distribution. Individual gene ρ phenotypes are shown in the top right, with black dots corresponding to individual contributing sgRNA ρ phenotypes. (B) Topology of γGI and τGI scores for *AUNIP*. As in Figure 4C. (C) Topology of γGI and τGI scores for *AUNIP* with enriched complexes indicated (Table S2; Methods). (D) Validation of interaction between *AUNIP* and BRCA1-A complex genes in K562 (top) and RPE1 (bottom) cell lines by targeted dual-sgRNA CRISPRi screen. Reference growth γ and drug sensitization ρ phenotypes were calculated for each treatment arm as in the full GI screen (Methods). Expected combinatorial phenotypes were determined by adding individual contributing phenotypes. Individual measurements from dual-sgRNA pairs contributing to each gene-level phenotype are shown by circles. (E) γGI, τGI, and νGI scores between *AUNIP*, the BRCA1-A complex, Fanconi anemia-associated genes, and genes within the *53BP1*-Shieldin-CST pathway. BRCA1-A complex and 53BP1-Shieldin-CST pathway genes were identified through literature review, while FA associated genes were determined by unbiased clustering performed in Figure 2E. ρ phenotypes for each gene are displayed below, with individual contributing sgRNA phenotypes shown by circles. (F) Model of antagonistic relationship between *AUNIP* and BRCA1-A complex operating in parallel to more canonical DNA repair genes. Gene names in gray were not included in our map.

To more systematically explore *AUNIP*-interacting genes, we identified protein complexes enriched among those genes (73 total; Figure 6C; Table S2; Methods). These enrichments confirmed relationships with FA pathway genes—some unmodified and some with greater synergism in presence of PARPi (9 νGIs)—and established connections with the ATR-ATRIP, 9-1-1 checkpoint, and BRCA1-A complexes. Interactions with the BRCA1-A complex, one of several cellular complexes formed with BRCA1, were of particular interest. *AUNIP* showed novel and possibly novel νGIs with most BRCA1-A complex genes targeted by our library (*i.e.*, buffering τGIs with no γGI), specifically *ABRA1* (*FAM175A*), *RAP80* (*UIMC1*), *BABAM1* (*MERIT40*), and *BRCC3* (Figures S6A and S6B). Notably, this trend did not hold for *BRCA1*, which, similar to other FA genes, instead demonstrated a negative τGI with *AUNIP* (Figure 6C).^117^ We next performed a set of focused, growth-based experiments in K562 and RPE1 cells. Resulting phenotypes, which were reproducible across replicates and related conditions (Figures S6C and S6D), confirmed that loss of BRCA1-A genes buffered PARPi sensitivity caused by loss of *AUNIP* (Figure 6D), even with a wider range of single-gene phenotypes observed in RPE1 cells.^118,119^ Additionally, further demonstrating context-dependence of gene function, perturbation of *AUNIP* alone sensitized cells to all three PARP inhibitors but not hydroxyurea, which causes replication fork stress through a different mechanism (Figure 6D).

A reasonable explanation for the buffering interactions between *AUNIP* and BRCA1-A genes is functional antagonism between promoting and suppressing repair at specific PARPi-induced DNA lesions. Although only minimally characterized, AUNIP has been shown to recruit CtIP to DNA lesions to promote DNA end resection,^120^ and the BRCA1-A complex has been separately reported to inhibit that process,^121,122^ with loss of complex genes resulting in an increased accumulation of CtIP-dependent nuclease activity at breaks. An antagonistic relationship between these genes could explain both phenotypes. We note that such relationships are canonically involved in regulating DNA repair processes. For example, as mentioned above, BRCA1 promotes DNA end resection,^123^ but when absent, gene products from the 53BP1-Shieldin-CST pathway prevent resection and potentiate the toxicity of PARP inhibitors.^3–12^ Accordingly, 53BP1-Shieldin-CST genes in our dataset^124–127^ demonstrated buffering τGIs and positive νGI scores with *BRCA1*, as well as FA pathway genes (Figure 6E). Examining GIs among these more canonical PARPi response factors, and with *AUNIP* and BRCA1-A genes, revealed three broad observations: first, 53BP1-Shieldin-CST genes minimally interacted with *AUNIP* in the νGI space, where interactions with *RIF1*, which appears to have secondary function in this context, and the upstream regulator *53BP1* were the primary νGI hits; second, positive νGI scores of BRCA1-A complex genes (excluding *BRCA1*) with *AUNIP* were generally stronger than those between the same BRCA1-A complex genes and the FA pathway set; third, while we observed many negative νGI hits between *AUNIP* and FA/BRCA1 genes (10 of 14) there were relatively few when looking at relationships of the latter set of genes with themselves (2 of 91). Altogether, these observations suggest that AUNIP and BRCA1-A control repair of specific PARPi-induced lesions similar to, but distinct from, more canonical PARPi response factors (Figure 6F).

## DISCUSSION

Here we present a large-scale analysis of differential genetic interactions in human cells. Using a DNA-repair focused CRISPRi sgRNA library targeting 147,153 gene pairs, we generated two static GI maps, a reference map from normally growing cells and one from cells exposed to niraparib. Next, we developed the analytical framework to compare these maps, thereby defining how gene-by-gene interactions change in the presence of PARPi-induced stress. These results reveal the complex landscape of coordinated gene functions required for genome stability maintenance normally and under one condition of genotoxicity. From this landscape, we reveal individual mechanistic insights and realize three broad achievements.

First, we show that measuring genetic interactions among similar genes can produce a high-resolution map of related processes. Specifically, despite biased representation of gene function within our library, we resolved functionally distinct gene modules, exploration of which then allowed inference of functional relationships at multiple levels; *i.e.*, between and within associated pathways and complexes. A clear example of this informational hierarchy is provided by our analysis of RAD51 accessory genes. Comprehensive inclusion of these genes within our map captured a remarkable level of within-module information, most predominantly functional redundancy between factors responsible for assembly, stabilization, and function of the RAD51 filament. Simultaneously, interactions between these genes and ones within a separate module (*i.e.*, BLM dissolvasome-associated genes) highlighted a general buffering relationship between more disparate, but still generally related, gene functions. Such informational density offers a rich resource for mechanistic exploration. To that end, we highlight that our data allows both the identification of specific gene-by-gene interactions *and* placement of those interactions within a broader context. For example, on their own, the intriguingly strong interactions between *RAD54L* and *PSMC3IP*-*MND1* warrant further study, but interactions between the latter and other RAD51 accessory genes also place those particular genes at the center of a process to which they have only recently been linked, somatic recombination.

Second, we developed a framework to evaluate differential genetic interaction networks in mammalian cells. Applying a categorization framework developed for yeast,^51^ we then cataloged context-specific roles for a host of genes after exposure to PARPi-induced cell stress. The utility of quantifying and categorizing these interactions at scale and across conditions is illustrated by the many context-specific interactions identified for *AUNIP*. This gene demonstrated relatively few interactions in our reference map, but together with a strong PARPi sensitization phenotype, the unusual number of *AUNIP* associated νGIs suggest a central role in managing PARPi-induced stress. Moreover, by exploring interactions between *AUNIP* and well characterized DNA repair genes, we proposed a model wherein AUNIP-promoted repair is suppressed by the BRCA1-A complex. Moving forward, we expect that our framework for mapping differential GIs will be applied across many conditions, within and outside of the field of DNA repair. Further illustrating the power of our chosen condition, though, the genetic network underlying PARPi response is itself context specific, with different gene-gene relationships expected in different genetic backgrounds. Applying our custom GI library tailored to the study of PARP inhibitors in different cell lines and cancer models will therefore also be instructive.

Third, we establish an important resource with clinical value. PARP inhibitors, including niraparib, are clinically approved drugs used to treat ovarian, breast, pancreatic, and prostate cancers. The identification of genes that modulate desired and undesired cellular activities of these drugs are thus of great interest. Our work provides a resource for exploring such genes across a wide range of genetic backgrounds; indeed, from one perspective, our differential GI map represents many parallel single-perturbation PARPi screens performed across 543 genetic backgrounds. Additionally, with respect to insights of potential clinical relevance, we found genetic interactions with *PARP2*. Hematological toxicities, such as thrombocytopenia, are some of the most common toxicities observed among patients treated with PARPi^128–131^ and are potentially exacerbated by PARP2 inhibition given its role in various hematopoietic processes.^132–135^ Our data identifies functional relationships with *PARP2* that support cell viability, thus providing key insight for future study of *PARP2* mechanisms.

In sum, we have shown that measuring differential genetic interactions at large-scale can provide a remarkably deep view of cellular processes that respond to changing stress environments. We expect that insights from the PARPi and genome stability interaction networks presented here will serve as a valuable resource for hypothesis generation and validation in future research. To aid such use of this resource, we have developed a web application for interactive exploration of the data (https://parpi.princeton.edu/map).

## ACKNOWLEDGEMENTS

We thank members of the Adamson lab for feedback and support throughout the course of this project, particularly P. Ravisankar, as well as the support of our friends and families. We thank L. Gilbert and B. Herken (University of California San Francisco), A. Ciccia (Columbia University), B. Engelhardt (Stanford University), and M. Levine (Princeton University) for helpful discussions. We thank W. Wang (Genomics Core Facility of Princeton University), and M. Schroeder (Princeton Research Computing). Research was supported by the National Institutes of Health (NIH) under award numbers R35GM138167 (B.A.), P30CA072720 (B.A.), and T32HG003284 (Princeton QCB training grant), as well as the Searle Scholars Program and Princeton University.

## AUTHOR CONTRIBUTIONS

Conception, D.S., J.L., and B.A.; Experimental methodology, D.S., J.L., and B.A.; Plasmid construction, J.L.; GI library design and cloning, D.S.; Screen and reagent optimization, D.S. and J.L.; Screens, J.L.; Validation, J.L. with assistance from Y.J.; Formal analysis, D.S.; Software and figure code, D.S.; Interpretation, D.S., J.L., and B.A.; Figures, D.S. and J.L.; Writing, D.S., J.L., and B.A. with input from Y.J.; Supervision, B.A.; Project administration, B.A.; Funding Acquisition, B.A.

## INCLUSION AND DIVERSITY

One or more of the authors of this paper self-identifies as a gender minority in their field of research. One or more of the authors of this paper self-identifies as a member of the LGBTQIA+ community.

## DECLARATION OF GENERATIVE AI AND AI-ASSISTED TECHNOLOGIES IN THE WRITING PROCESS

During the preparation of this work the authors used ChatGPT to reduce the word count in the summary. After using this tool, the authors reviewed and edited the content as needed and take full responsibility for the content of the publication.

## METHODS

### RESOURCE AVAILABILITY

#### Lead contact

Further information and requests for resources and reagents should be directed to and will be fulfilled by the Lead Contact, Britt Adamson.

#### Material availability

Plasmids and dual-sgRNA CRISPRi libraries (lDS001 and lYJ001) generated in this study will be deposited to Addgene.

#### Data and code availability

Processed data from primary genome-scale screen and genetic interaction screen are available as supplementary tables to this manuscript. Raw sequencing data from all screens will be deposited to the NCBI GEO repository. Scripts used to process data from the GI screen using R, as well as to reproduce manuscript figures, will be available on Github at https://github.com/simpsondl/PARPi_interactions. Data from both the reference and PARPi genetic interaction screens can also be accessed interactively through our accompanying web application located at https://parpi.princeton.edu/map.

### EXPERIMENTAL MODEL AND SUBJECT DETAILS

Cell lines used in this study were Lenti-X 293T, K562 CRISPRi, and RPE1 CRISPRi cell lines. K562 CRISPRi and RPE1 CRISPRi cells expressing dCas9-BFP-KRAB (pHR-SFFV-dCas9-BFP-KRAB; Addgene, 46911) were described previously.^136^ Lenti-X 293T cells were purchased from Takara (#632180). K562 CRISPRi and RPE1 CRISPRi cell lines were authenticated by analysis of short tandem repeats as exact matches to the corresponding line, CCL-243 (K562, female) and CRL-4000 (hTERT RPE1, female) from ATCC. Lenti-X 293T cell lines were authenticated by analysis of short tandem repeats as a similar match to the corresponding line CRL-3216 (293T) from ATCC. Cells tested negative for micoplasma. K562 CRISPRi cells were grown in RPMI 1640 with L-glutamine and 25 mM HEPES (Corning or Gibco) supplemented with 10% FBS (Gibco #10437-028), 100 U/mL penicillin, 100 μg/mL streptomycin, and 0.292 mg/mL L-glutamine. 293T cell lines were grown in DMEM with 4.5 g/L glucose and sodium pyruvate without L-glutamine (Corning) supplemented with 10% FBS (Gibco #10437-028), 100 U/mL penicillin and 100 μg/mL streptomycin. RPE1 CRISPRi cells were grown in DMEM/F12 with L-glutamine and 15 mM HEPES (GIBCO) supplemented with 10% FBS (Gibco #10437-028), 100 U/mL penicillin and 100 μg/mL streptomycin. Cells were grown at 37°C with 5% CO_2_ in standard tissue culture incubators. Niraparib, veliparib, talazoparib, and HU were dissolved in DMSO.

### METHOD DETAILS

#### Plasmid construction

Our dual-sgRNA CRISPRi library was built based on two sgRNA expression vectors (pJL051 and pJL052) which were modified from published sgRNA lentiviral plasmids (pLG_GI2 and pLG_GI3).^48^ Modifications were made by replacing the sequence between restriction sites AvrII and NsiI in pLG_GI2 with intended restriction sites (NsiI and AscI) and by replacing the sgRNA cassette between XhoI and BamHI in pLG_GI3 with a new cassette containing a different sgRNA constant region and the intended restriction sites (SbfI and AscI) at the 3′ end. The resulting sgRNA expressing cassettes have a modified mouse U6 promoter (pJL051) or a modified human U6 promoter (pJL052) followed by a sgRNA protospacer and a constant region CR1 (pJL051) or CR3 (pJL052; Figure S1B).^137^ Both vectors have BlpI and BstXI restriction sites flanking the sgRNA sequence for cloning. The vectors also co-express the puromycin resistant gene and BFP from an EF-1α promoter.

For the arrayed validation experiments, individual sgRNAs were delivered using pU6-sgRNA EF1Alpha-puro-T2A-GFP (Addgene, 111596)^48^ or pU6-sgRNA EF1Alpha-puro-T2A-BFP (Addgene, 60955).^55^

#### Virus preparation

Lentivirus was produced in Lenti-X 293T cells by co-transfection of transfer plasmids (single or library) and packaging plasmids for expression of HIV-1 gag/pol, rev, and VSV-G envelope protein using TransIT®-LT1 Transfection Reagent (Mirus) with ViralBoost Reagent (Alstem, Inc.). Virus-containing supernatants were collected and stored frozen. Viral titers were determined by testing transductions prior to screening.

#### Genome-scale PARPi screen

K562 CRISPRi cells were transduced with the hCRISPRi-v2 top 5 library (Addgene #83969)^138^ with 840X representation. Transduction was supplemented with 8 μg/mL polybrene and conducted in many wells of multiple 6 well plates with centrifugation (∼2 hours at 1000 x g). After centrifugation, cells were pooled and resuspended in fresh, complete RPMI to ∼0.5e6 cells per mL and maintained in an Multitron Incubation Shaker (Infors HT) at appropriate speeds throughout the screen. 3 days post transduction, cells were 25% BFP+ (sgRNA library marker) and were selected with 1 μg / mL puromycin until selection was complete. On day 10 post transduction, treatment started (T0). T0 samples were collected and remaining cells were split into four populations with two dosed with niraparib (4.5μM, ∼IC30) and two treated with vehicle (DMSO). Cells were typically maintained at densities between ∼0.5e6 and ∼1e6 cells per mL (splitting as necessary). A 2000X representation was maintained for each replicate during the treatment. 13 days post transduction, cells were spun down and resuspended in drug-free media. 19 days post transduction, cells were collected (endpoint) and cell pellets were processed to generate sequencing libraries as follows. Genomic DNA was extracted using the NucleoSpin® Blood XL, Maxi kit for DNA from blood (Macherey-Nagel). The sgRNA loci were amplified with an indexing 5′ primer (5′-aatgatacggcgaccaccgagatctacacgatcggaagagcacacgtctgaactccagtcacNNNNNN gcacaaaaggaaactcaccct) and a common 3′ primer (5′-CAAGCAGAAGACGGCATACGAGATCGACTCGGTGCCACTTTTTC). Each reaction (100 μL total volume) contained 10 μg of genomic DNA (measured by Nanodrop) and 1 μM of each primer and was run on a thermocycler with the following program: 1 cycle of 30 seconds at 98°C; 22 cycles of 10 seconds at 98°C, followed by 75 seconds at 65°C; 1 cycle of 5 minute at 65°C; 4°C hold. Between 410 to 570 reactions were set up for each arm to obtain 1800X coverage. The resulting products were purified for sequencing using SPRIselect Reagent (Beckman Coulter) in a double-sided 0.65X-1X reaction followed by 1X reaction twice. Indexed samples were pooled in equal molar ratio prior to sequencing.

#### Design of dual-sgRNA CRISPRi library

A total of 543 genes were included in our dual-sgRNA CRISPRi library (Table S1). To choose genes for our map with roles in DNA repair and associations with PARP inhibitor response, we used a combination of results from our genome-scale niraparib screen, a thorough literature review, and supplementary pathway enrichments. As the first step in our literature review, we collected data from 24 previously published single-perturbation CRISPR screens reported to identify genes that confer sensitization or resistance to PARP inhibition.^5,6,8,60–63^ These screens used different PARP inhibitors, cell lines, perturbation strategies, and screening protocols, but evaluation of the reported data with a consensus approach identified a high-priority set of 177 genes, defined as those that identified as hits in at least three screens. Concurrently, we performed our own single-perturbation, genome-scale CRISPRi screen using an understudied PARPi, niraparib. Using data from this screen, we selected 247 genes (Table S1), 68 of which were also identified by the literature review. As the second step in our literature review, we identified and supplemented our selected genes with a further 62 genes known or suspected to be involved in PARPi response. For example, *PARP2* does not have a PARPi sensitization phenotype and as such was not identified by either our screen or review of previous screens (Figure 1B). However, the gene is one of two molecular targets of niraparib, and we therefore included it in our library at this stage. Finally, we completed our gene set with a selection of 125 genes annotated to top pathways enriched among the previously identified set of 418 genes using Reactome pathway database (Table S2).

For each gene, we chose sgRNAs with the largest magnitude of phenotype and consistent sign (*i.e.*, both sensitizing or resistance) observed in our genome-scale screen. Our final library included 1086 gene-targeting sgRNAs (2 for each gene) and an additional 55 non-targeting controls for a total of 1141 sgRNAs (Table S1). All spacer sequences were of the form G[N19]. Among sgRNAs included in the library, reference growth phenotypes (γ) ranged from -0.45 to 0.06 in the genome-scale screen, and growth phenotypes measured with PARPi treatment (τ) were -0.66 to 0.26 (Figure S1). Due to the cloning process used, each of the 1141 guides could appear in either position in our library vector, resulting in up to 1,301,881 (1141 x 1141) distinct “construct” combinations, representing 147,153 (543 choose 2) gene pairs.

#### Cloning of dual-sgRNA CRISPRi library

sgRNA targeting sequences were ordered as a Twist oligo pool containing two subpools with different previously verified primer binding regions^55^ and different constant region fragments for cloning into the pJL051 and pJL052 vector backbones. Each subpool was structured similarly with primer amplification binding regions on the 5′ (5′-ATTTTGCCCCTGGTTCTT for pJL051; 5′-TCACAACTACACCAGAAG for pJL052) and 3′ (5′-CCAGTTCATTTCTTAGGG for pJL051; 5′-GCAACACTTTGACGAAGA for pJL052) ends and BstXI (5′-CCACCTTGTTG) and BlpI (5′-GTTTAAGAGCTAAGC for pJL051; 5′-GTTTCAGAGCTAAGC for pJL052) restriction enzyme sites flanking the sgRNA targeting sequence. Twist oligo subpools were PCR amplified using 2X Phusion Master Mix, 0.5μM forward primer (5′-ATTTTGCCCCTGGTTCTTCCAC for pJL051; 5′-TCACAACTACACCAGAAGCCAC for pJL052), 0.5μM reverse primer (5′-CCCTAAGAAATGAACTGGGCTT for pJL051; 5′-TCTTCGTCAAAGTGTTGCGCTT for pJL052), and 0.1pmol resuspended Twist oligo pool with the following conditions: 1 cycle of 30 seconds at 98°C; 15 cycles of 15 seconds at 98°C, followed by 15 seconds at 56°C, followed by 15 seconds at 72°C; 1 cycle of 10 minutes at 72°C; 7°C hold. Two PCR reactions were run for each subpool, then PCR products were aggregated and purified using Machery-Nagel NucleoSpin Gel and PCR Clean-up kit and quantified using Nanodrop. Each of our two vector backbones (pJL051 and pJL052) and the two amplified oligo pools were subject to a BstXI-BlpI double digest, then the fragments were gel extracted and ligated at room temperature for 15 minutes using T4. Ligation products were electroporated using Mega-X cells resulting in an averaged >1000 transformants for each library element. The sgRNA loci were amplified from plasmid libraries for validation with an indexing 5′ primer (5′-AATGATACGGCGACCACCGAGATCTACACGATCGGAAGAGCACACGTCTGAACTCCAGTCACGCCAATGCACAAAAGGAAACTCACCCT for pJL051; 5′-AATGATACGGCGACCACCGAGATCTACACGATCGGAAGAGCACACGTCTGAACTCCAGTCACATCACGGGACTATCATATGCTTACCGTAAC for pJL052) with accompanying 3′ primers (5′-CAAGCAGAAGACGGCATACGAGATCGACTCGGTGCCACTTTTTC for pJL051; 5′-CAAGCAGAAGACGGCATACGAGATGGCGGTAATACGGTTATCCA for pJL052). Intermediate sgRNA libraries were sequenced on Illumina Miseq at >450X coverage (see “Sequencing”) and confirmed that 90% of library elements are within ∼3x coverage of each other (95th percentile coverage/5th percentile coverage = 3.14 for pJL051; 95/5 = 3.62 for pJL052) for both libraries.

Intermediate sgRNA libraries were then combined to form the dual-sgRNA CRISPRi library lDS001. pJL051 was digested using NsiI and AscI, while pJL052 was digested using SbfI and AscI, as these form isocaudamer pairs resulting in overhangs that can be ligated together. The digested libraries were gel-extracted and ligated at room temperature for 15 minutes using T4. Multiple T4 ligations (32) were combined and then cleaned of excess salts using Machery-Nagel NucleoSpin Gel and PCR Clean-up kit to generate an adequate concentration of ligated DNA. The resulting vector contains the individual library cloned into pJL051 in the first position (position A) and the individual library cloned into pJL052 in the second position (position B). This concentrated ligation product was electroporated with Lucigen Endura electrocompetent cells, which we found increased electroporation efficiencies by 6-fold over Mega-X electrocompetent cells for the lentivirus backbones used in this screen, and resulted in >100 transformants for each library element. Cultures were grown at scale to generate large amounts of the final dual-sgRNA library which was validated using an Illumina Novaseq 6000 at >100x coverage with the sequencing approach outlined in Figure S1C (see “Sequencing”). Sequencing confirmed the presence of 99.96% of library elements in the cloned plasmid library (1,301,379 constructs), with individual positions showing a similar distribution of sgRNAs to that of the corresponding sub-libraries (95/5 = 3.17 for position A; 95/5 = 3.08 for position B). Additionally, the Gini inequality coefficient, which measures the degree of inequality in a frequency distribution on a scale of 0 (all frequencies equal) to 1 (all frequencies equal to 0 except one), showed a relatively low degree of inequality in our constructed dual-sgRNA CRISPRi library (Figure S1D), backed by a 95/5 ratio of 5.79.

For the pooled validation experiments in Figure 6, individual sgRNAs were first cloned into pJL051 and pJL052. Constructs with selected sgRNAs were then pooled to create arrayed sub-libraries. pJL052 sub-libraries were digested with SbfI and AscI and the extracted fragments were ligated into corresponding pJL051 sub-libraries digested with NsiI and AscI to form intended sgRNA combinations. Ligation products were transformed into Stbl3 chemically competent cells and isolated using column-based purification. The resulting dual-sgRNA sub-libraries were pooled in equal amounts for each sgRNA pair and validated by Illumina sequencing.

#### Large-scale interaction screen

K562 CRISPRi cells were transduced with the dual-sgRNA CRISPRi lentiviral library with a 350X representation per construct at a low MOI (∼0.25) as described above. Two biological replicates were performed. Cells were maintained at 0.5e6 to 0.6e6 per mL in the multitron incubator and were selected with 3 μg/mL puromycin 2 days post transduction. 9 days post transduction, treatment started (T0). T0 samples were collected (500X representation) from each replicate and the remaining cells of each replicate were divided into two populations, one of which was dosed with niraparib (2 μM, roughly the LC30 determined for cells transduced with the interaction sgRNA library) and the other with vehicle (DMSO). A 950X representation was maintained for each arm throughout the treatment. Cells were maintained at densities between 0.5-0.6e6 cells per mL during the treatment (splitting as necessary). Treatment lasted for 3 days, after which cells were spun down, resuspended in drug-free media and were allowed to grow for an additional 6 days (Figure S1E). 18 days post transduction, cells were collected (500X representation) from the niraparib-treated arms (PARPi endpoint samples) and DMSO-treated arms (reference endpoint samples), respectively. Cell pellets were processed to generate sequencing libraries as follows. Genomic DNA was extracted using the NucleoSpin® Blood XL, Maxi kit for DNA from blood (Macherey-Nagel) and treated by DNase-free RNase A followed by ethanol precipitation. The sgRNA loci were amplified with an indexing 5′ primer (5′-AATGATACGGCGACCACCGAGATCTACACNNNNNNNNCAGCACAAAAGGAAACTCACC) and an indexing 3′ primer (5′-AAGCAGAAGACGGCATACGAGATGTGACTGGAGTTCAGACGTGTGCTCTTCCGATCTNNNNNNNNGGCGGTAATACGGTTATCCA). Each reaction (100 μL total volume) contained 10 μg of genomic DNA (measured on Invitrogen Qubit 4 Fluorometer) and 1 μM of each primer and was run on the Bio-Rad C1000 thermal cycler with the following program: 1 cycle of 30 seconds at 98 °C; 20 cycles of 10 seconds at 98 °C, followed by 75 seconds at 65 °C; 1 cycle of 5 minute at 65 °C; 4 °C hold. At least 450X coverage was maintained for each sample during PCR amplification. Aliquots of the PCR products were purified for sequencing using SPRIselect Reagent (Beckman Coulter) in two double-sided 0.45X-0.8X reactions and two 0.8X reactions. Indexed samples were pooled in equal molar ratio prior to sequencing.

#### Generation and validation of knock-out cell lines

spCas9 from Integrated DNA Technologies (Alt-R® spCas9 Nuclease V3, 1081059) were complexed with *PARP2* single guide RNAs from IDT (predesigned Alt-R® CRISPR-Cas9 crRNA Hs.Cas9.PARP2.1.AA and Alt-R® CRISPR-Cas9 tracrRNA). 2.5e5 RPE1 CRISPRi cells were nucleofected with 3μM RNP complex using an SE Cell Line 4D X Kit S (Lonza Bioscience) on a 4D-Nucleofector (Lonza Bioscience) according to manufacturer’s instructions (program EA-104). 3 days post electroporation, single cells were sorted into 96-well plates using FACS cell sorter (BD Biosciences FACSAria Fusion). Single cell clones were grown and expanded for 3 to 4 weeks before being frozen down. Genomic DNA from each clone was extracted and the *PARP2* locus was amplified with primers oJL392 (5′-TGGAGTTCAGACGTGTGCTCTTCCGATCTGGTCTAAGGAAAGACCAGG) and oJL393 (5′-ACACTCTTTCCCTACACGACGCTCTTCCGATCTGGATGCTCTCTGAGATATCC). The PCR products of individual clones were indexed (with primers 5′-CAAGCAGAAGACGGCATACGAGATNNNNNNNNGTGACTGGAGTTCAGACGTGTGCTCTTC and 5′-AATGATACGGCGACCACCGAGATCTACACNNNNNNNNACACTCTTTCCCTACACGAC), pooled, and sequenced. The sequencing results were analyzed using CRISPResso2.^139^

#### Arrayed validation experiments

*Performed with CRISPRi.* Two single-perturbation sgRNA vectors targeting different genes marked with either GFP or BFP were co-transduced into K562 CRISPRi or RPE1 CRISPRi cells. Four populations of the transduced cells were detected on a flow cytometer: BFP-GFP-(uninfected), GFP+ BFP-, BFP+ GFP-, and BFP+ GFP+. Starting from 6 days post transduction, cells were recorded for the percentage of the four populations by flow cytometer every 2 or 3 days until day 15 or day 21. The change in percentage of each population over time was normalized to the uninfected population and converted to log_2_ fold change. Expected dual-sgRNA phenotypes were obtained by adding two single-sgRNA phenotypes. Triple replicates were performed for each experiment. Interaction effect sizes used in Figure S3D were calculated using Cohen’s D, or standardized mean difference, between the distributions of expected and observed combinatorial phenotypes, implemented in R package effsize.^140^ *Performed with knockout cell lines.* Individual sgRNAs targeting *RNASEH2C* or non-targeting sgRNAs marked with BFP were transduced into WT or PARP2 KO RPE1 CRISPRi cells in triplicate. 9 days after transduction, each replicate was split into 2 arms, with one treated with niraparib (1 μM) and the other with vehicle (DMSO), and the percentage of BFP+ and BFP-populations in each sample was recorded by flow cytometer every 3 days until day 21. The change in percentage of each population over time was normalized to the uninfected population and converted to log_2_ fold change.

#### Pooled validation experiments

K562 CRISPRi or RPE1 CRISPRi cells were transduced with the focused sgRNA library at ∼0.2 MOI and were selected with puromycin starting 2 or 3 days post transduction (3 μg/mL for K562, 8 μg/mL for RPE1). 9 days post transduction, T0 samples were collected and remaining cells were aliquoted and treated with drugs or vehicle (DMSO) in triple replicates. The drug dosage were 1 μM, 5 nM, 4 μM, and 100 μM for niraparib, talazoparib, veliparib, and HU, respectively, in RPE1 CRISPRi cells and 2 μM, 10 nM, 9 μM, and 25 μM for niraparib, talazoparib, veliparib, and HU, respectively, in K562 CRISPRi cells. 3 days later, media was replaced with drug-free media and cells were allowed to grow for an additional 6 days, except for HU-treated K562 CRISPRi samples where cells were dosed with 100 μM HU again. 18 days post transduction, cells were collected and cell pellets were processed to generate sequencing libraries as described for the large-scale interaction screen, except for the following differences: RNase A treatment was done during cell lysis before genomic DNA was extracted; each 100 μL PCR reaction contained about 1 μg or 2 μg of genomic DNA and PCRs were run for 22 cycles; PCR products were purified using SPRI in one double-sided 0.45X-0.8X reaction and three 0.8X reactions.

#### Quantitative RT-PCR

Cells were transduced with single sgRNA vectors as described above, selected by puromycin, and harvested 6 days post transduction. Total RNA was isolated using the Quick-RNA Miniprep Kit (Zymo Research). Quantitative RT-PCR reactions were assembled using the Power SYBR Green RNA-to-CT 1-Step Kit (Invitrogen) according to the manufacturer’s instructions. Reactions were run on a ViiA 7 real-time PCR system (Thermo Fisher). Primer sequences are listed in Table S7.

#### Sequencing

*Single-perturbation genome-scale screen.* Sequencing was performed on an Illumina NovaSeq 6000 System with 20% phiX spike-in with single-end reads: I1 = 6 nt, sample index; R1 = 50 nt, sgRNA identity. Custom primer was used for the R1 read (Table S7). *Validation of dual-sgRNA position-specific sgRNA libraries.* Individual component libraries cloned into pJL051 and pJL052 backbones were validated using an Illumina MiSeq system with 10% phiX spike-in with a single-end read: I1 = 8 nt, sample index; R1 = 50 nt, sgRNA identity. Custom primer was used for R1 read (Table S7). *dual-sgRNA library validation and interaction screen*. Library of the large-scale interaction screen was sequenced on an Illumina NovaSeq 6000 System with a 15%-25% phiX spike-in with paired end reads: I1 = 40 nt, sgRNA at position B; I2 = 8 nt, sample index; R1 = 40 nt, sgRNA at position A; R2 = 8 nt, sample index (Figure S1C). Custom primers were used for I1, I2, and R1 reads, as well as for phiX spike-in (Table S7). *Pooled validation experiments.* Sequencings were performed on an Illumina MiSeq system with 25% phiX spike-in with paired-end reads: I1 = 40 nt; I2 = 8 nt; R1 = 40 nt; R2 = 8 nt. Same custom primers as the interaction screen were used for I1 and R1 reads. *Validation of knock-out cell lines*. Sequencing of PARP2 knock-out clones was performed on an Illumina MiSeq System with 10% phiX spike-in with single-end reads: I1 = 8 nt; I2 = 8 nt; R1 = 300 nt.

#### Flow cytometry

Flow cytometry data was collected with an Attune NxT Flow Cytometer (Thermo Fisher).

### STATISTICAL ANALYSIS

#### Sequence alignment and count generation

*Single-perturbation genome-scale library.* Reads were aligned to a reference index using BWA,^141^ those with mapping quality ≤ 10 were filtered, and counts were determined for each sgRNA using the surviving reads (Table S1). *dual-sgRNA libraries.* To ensure high quality read assignment, we used a two-step aligning and filtering process. Reads consist of 19 nt corresponding to an sgRNA target sequence followed by 21 nt from the associated sgRNA constant region. First, we analyzed the constant region in each read using bowtie2^142^ to soft trim 19 nt from the 5′ end and map reads to reference constant regions. Reads which mapped constant sequences in the incorrect orientation or which had more than 1 mismatch were discarded using the Picard FilterSamReads module. We reasoned that these reads represented inactive sgRNAs with defects from cloning. sgRNA target regions from the remaining reads were then analyzed by using bowtie2 to soft trim 21 nt from the 3′ end, isolating only the first 19 nt, and align the remaining read fragments to an sgRNA reference library. Reads with mapping quality ≤ 5 were filtered, and read counts for each position-specific sgRNA pair, or “construct” (denoted by X/Y), were then determined from the remaining aligned reads.

#### Single-perturbation, genome-scale library analysis

Screen data was analyzed using a consensus approach, combining outputs from multiple tools to derive a set of hits. Using an established methodology,^138^ we first calculated growth phenotypes in reference (gamma, γ) and PARPi-treated (tau, τ) conditions as well as PARPi sensitization (rho, ρ) phenotypes for each sgRNA as follows: a pseudocount of 10 was added to all counts (see “Sequence alignment and count generation”), then enrichment values were calculated for each phenotype, followed by normalization to non-targeting controls and the number of population doublings. Specifically, enrichments were determined as γ = log2(fraction counts at DMSO endpoint / fraction counts at T0); τ = log2(fraction counts at PARPi endpoint / fraction counts at T0); and ρ = log2(fraction counts at PARPi endpoint / fraction counts at DMSO endpoint), then, for each phenotype, the median enrichment among non-targeting sgRNAs was subtracted from all enrichments, and these values were then divided by the number of population doublings in each endpoint (reference R1 = 7.65; reference R2 = 7.76; PARPi R1 = 4.71; PARPi R2 = 6.07; ρ R1 = 7.65 - 4.71 = 2.94; ρ R2 = 7.76 - 6.07 = 1.69). These phenotypes were supplied to CRISPhieRmix,^143^ and counts were supplied to each of Model-based Analysis of Genome-wide CRISPR-Cas9 Knockout (MAGeCK),^144^ DrugZ,^145^ and the ScreenProcessing pipeline^138^ to analyze results from the genome-scale screen. Tools were run under default parameters. Genes which were identified in a hit using two or more tools (247) were included in the interaction library. Gene-level significances produced by MAGeCK through robust rank aggregation^146^ and phenotypes calculated via the described approach are used in Figure 1B and are available in Table S1.

#### Calculating construct and sgRNA phenotypes

Because each gene in our library is represented by two sgRNAs and each sgRNA can maximally occur in either position in the library vector (A or B), there are up to 8 possible dual-sgRNA “constructs” associated with each pair of genes in our library. For each of these constructs, we calculated growth phenotypes in reference (gamma, γ) and PARPi-treated (tau, τ) conditions in addition to PARPi sensitization phenotypes (rho, ρ). First, we identified and removed constructs containing individual sgRNAs represented at low frequency. Specifically, we calculated the median coverage for each sgRNA across replicates in each position and filtered constructs with any sgRNAs demonstrating a median coverage <30 reads in either position at T0. We then also removed any dual-sgRNA constructs with <25 reads in either replicate at T0, and to further avoid effects from low frequency representation, we added a pseudocount of 10 to all surviving construct read counts. Next, we defined an enrichment value for individual constructs as log_2_ of the fraction of reads (*i.e.*, reference or PARPi) in endpoint samples divided by the fraction of reads for that construct at T0 (Figure S2A). Enrichments were normalized to non-targeting controls, for γ and τ separately, by subtracting the median enrichment of non-targeting sgRNA pairs (NT/NT) and adjusted by dividing the total number of population doublings that occurred from T0 until endpoint (reference R1 = 7.19, reference R2 = 7.73, PARPi R1 = 5.32, PARPi R2 = 5.36). The resulting values constitute our orientation-dependent γ and τ “construct phenotypes”. To calculate construct ρ phenotypes, we divided the fraction of reads at the PARPi endpoint by the fraction of reads at the reference endpoint, normalized to non-targeting controls, and then adjusted the resulting values by dividing by the difference in the number of population doublings observed in the PARPi-treated and reference arms (R1 = 1.87, R2 = 2.37). To determine individual “sgRNA phenotypes”, we averaged the phenotypes of all constructs wherein a given sgRNA was paired with a non-targeting sgRNA, independent of orientation (*i.e.*, including X/NT and NT/X) but separately for each replicate. Similarly, “single-gene phenotypes” were defined as the average of all sgRNA phenotypes for sgRNAs targeting the same gene.

#### Calculating model-specific, sgRNA-level GI scores

We calculated γGI and τGI scores between sgRNAs using a model-based approach essentially as previously described.^48^ γGI and τGI scores were calculated independently using condition-specific data (reference or PARPi) and the following procedure: to reduce noise and increase sensitivity, we first calculated “sgRNA pair phenotypes” (γ and τ) for each unique sgRNA pair by averaging construct phenotypes (see “Calculating construct and sgRNA phenotypes”) with the same sgRNAs in different orientations (X/Y and Y/X), and then averaging sgRNA pair phenotypes across replicates. This process collapsed data from orientation-specific constructs into orientation-independent values for the vast majority of sgRNA pairs (99%) but, due to filtering, retained information from only one construct (X/Y or Y/X) for some sgRNA pairs (1%). We then determined the Pearson correlation between sgRNA pair phenotypes associated with a single “query” sgRNA and the corresponding sgRNA phenotypes for each partner or “object” sgRNA (see “Calculating construct and sgRNA phenotypes”). Note that for this step, we pulled phenotype information from an uncollapsed dataset where sgRNA pair phenotypes were recorded using construct identifiers and thus most measurements (99%) were listed twice. We accessed phenotypes non-redundantly by pulling sgRNAs according to their position in the identifier, defining position B as the “query position” and position A the “object position”. To ensure high-quality interaction scores, we removed information associated with queried sgRNAs for which Pearson *r* < 0.2. Then, for each remaining query sgRNA, we fit a quadratic model to sgRNA-pair phenotypes associated with that query and corresponding sgRNA phenotypes from object sgRNAs. We defined model-specific “sgRNA-level GI scores” as the difference between observed sgRNA-pair phenotypes and model-defined combinatorial phenotypes (*i.e.*, the model residuals), normalized for each query by the standard deviation of interaction scores from non-targeting object sgRNAs. Through this methodology, each individual sgRNA pair phenotype with two contributing construct phenotypes (X/Y and Y/X) generated two different, model-specific, sgRNA-level GI scores, one from the model determined by query X and one from query Y, while sgRNA pair phenotypes with only one contributing construct phenotype (Y/X or X/Y) generated one sgRNA-level GI score, from either the model determined by query X or the one from query Y.

We found that calculating PARPi-specific interaction scores in an analogous model-based way using ρ phenotypes resulted in high variance scores called from models with poorer explanatory power than observed for γ and τ phenotypes. Specifically, while ∼35 to 40% of sgRNA models had an adjusted R^2^ value less 0.8 when using γ or τ phenotypes, 85% of sgRNA models built from ρ phenotypes had adjusted R^2^ values in this range. We also observed that sgRNAs with strong individual ρ phenotypes (defined as |ρ| > 0.3 for this analysis) had an increased frequency of poor R^2^ values (adjusted R^2^ < 0.8; Fisher’s exact p = 1.57e-55), indicating that those genes in which we are most interested, those with a direct role in responding to PARPi-induced damage, are more likely to generate unreliable interaction scores with this approach. To overcome this limitation, we defined a differential interaction score, termed a νGI score, for each sgRNA pair using its associated γGI and τGI scores. To calculate νGI scores, sgRNA-level GI scores from both the γ and τ settings were unified by keeping only those called from equivalent models in both conditions, and then normalized γGI scores were subtracted from the corresponding model-specific, normalized τGI scores.

#### Calculating gene-level GI scores

Gene-level γGI, τGI, and νGI scores were calculated as the average of all model-specific, sgRNA-level GI scores associated with a given gene pair in each setting (see “Calculating model-specific, sgRNA-level GI scores”). Because our library typically had two sgRNAs per gene post-filtering, most gene pairs therefore had four sgRNA pairs, which together contributed eight model-specific, sgRNA-level GI scores per condition. However, as the query sgRNA correlation filter (see “Calculating model-specific, sgRNA-level GI scores”) was applied to γ and τ phenotypes independently to preserve as much condition-specific information as possible, some sgRNA-level GI scores appear only in one context and not the other. When calculating gene-level νGI scores, such sgRNA-level scores are excluded, which can cause a discrepancy when comparing νGI scores to the underlying γGI and τGI scores. We have annotated which sgRNAs were modeled in each condition in Table S1 and noted gene pairs with such a difference in Table S3 with the interaction category “uncharacterized”. For sgRNA pairs targeting the same gene, only sgRNA-level GI scores from distinct sgRNA combinations were used to define a “single-gene GI score” (*i.e.*, X-1/X-1 is excluded when determining single-gene GI scores for gene X). Additionally, to generate pseudogene controls, we randomly assigned the 55 non-targeting sgRNAs within our library to 29 pseudogenes. Similar to true gene pairs post-filtering, 10% of these pseudogenes were supported by one sgRNA and 90% were supported by two. sgRNA-level GI scores involving one or two non-targeting sgRNAs were averaged according to these pseudogenes to produce a distribution of gene-level interactions involving non-targeting sgRNAs.

As poorly represented elements (*i.e.*, sgRNAs and constructs) were filtered from the data, we considered the effects of such dropout on gene-level GI scores. For clarity, when one of 4 possible sgRNAs targeting a pair of genes was filtered from the dataset, the maximum number of contributing model-specific, sgRNA-level GI scores dropped from 8 to 4, and if 2 such sgRNAs were filtered (*i.e.*, one from each gene), then at most 2 such scores were used. Gene-level interaction scores “supported” by fewer model-specific, sgRNA-level scores had increased variance and identified higher proportions of interactions as significant, with scores supported by ≤ 3 being particularly susceptible to this effect (Figure S3E). When considering individual genetic interactions from our dataset, the number of supporting scores underlying each gene-level measurement should therefore be considered (Table S3).

#### Assigning significance to GI scores

To estimate statistical significance of γGI and τGI scores, we used permutation testing, a non-parametric approach that uses the data itself to estimate the null distribution rather than making assumptions about the underlying distribution. This strategy is well-suited for our use case where standard probability models (*e.g.*, Gaussian) poorly fit the distribution of observed GI scores. Sets of GI scores from null distributions were generated by permutation testing in the following way: we shuffled labels associated with construct phenotypes and recalculated individual sgRNA phenotypes (see “Calculating construct and sgRNA phenotypes”), rebuilt sgRNA-level models (performing no filtering of query sgRNAs) and, using those models, determined new gene-level γGI and τGI scores (147,658 for each; see “Calculating model-specific, sgRNA-level GI scores” and “Calculating gene-level GI scores”). This process was repeated 2,000 times for γGIs and for τGIs to generate a set of 295,316,000 (147,658 x 2,000) gene-level GI scores estimating each null distribution. For each gene pair, we then calculated a two-sided p-value as the number of scores from the interaction screen or from null distributions at or exceeding the magnitude of the observed score, divided by the total number of scores from the original and permuted data. To control the false discovery rate (FDR), q-values and local FDR were determined for each set of p-values (Table S3; qvalue package^147^ in R). While q-values control the overall FDR of a given set of p-values, local FDR defines the likelihood of individual measurements being false discoveries. We define “high-confidence interactions” as those with local FDR ≤ 0.01 in the reference and PARPi conditions, and “significant interactions” as those with q-value ≤ 0.05. Similarly, to estimate the null distribution for νGI scores, labels for the unified set of observed model-specific sgRNA pair τGI scores were shuffled independently of those for γGI and νGI scores were then recomputed, generating 287,758,000 (143,879 x 2,000) gene pair scores from null distributions (see “Calculating model-specific, sgRNA-level GI scores” and “Calculating gene-level GI scores”). Two-sided p-values, q-values, and local FDR values for observed νGI scores were calculated, and νGI “hits” were defined using a threshold of q-value <= 0.2 (Table S3). Increased tolerance for false discovery in the ν space was intended to account for the increase in noise observed by normalizing measurements across conditions.

#### Gene annotations

Genes selected for our dual-sgRNA library were supplied to DAVID Functional Annotation Tool^148,149^ for functional annotation using the UniProt Keyword “Biological Process”,^150^ Gene Ontology (GO) “Biological Process”,^151,152^ and KEGG Pathway^153,154^ databases. Annotation categories were clustered by DAVID 2021 and the top clusters were used to define general annotation categories (Figure 1C; Table S2). Genes annotated to *DNA damage repair* were re-queried against the same databases to provide more specific annotations (Table S2). Reactome pathway database^64^ was queried separately to identify enriched pathways among GI library genes. Previously validated interactions used in Figures 2D and 3B were identified using STRING v11.5^88^ by searching for “experimentally validated interactions” or “gene fusions” with medium (score ≥ 0.4), high (score ≥ 0.7), and highest confidence (score ≥ 0.9). Note that in Figure 2D, the set of 1,950 experimentally validated interactions identified by STRING only partially overlapped with interactions identified as significant in our dataset (263 of 1950, 13.5%). This may partly be explained by two factors: (1) the previously validated interactions identified by STRING are physical interactions rather than genetic, as with *FANCA* and *FANCB*, and (2) not all genetic interactions will show a phenotype in all contexts. Finally, PANTHER^155,156^ was used to query both the Reactome pathway and GO Cellular Component databases for analyses related to Figures S3F and 6C. For analysis in Figure 6C, genes were selected by having either a νGI hit (FDR ≤ 0.20) or high-confidence τGI (FDR ≤ 0.05) with *AUNIP*.

#### Clustering and visualization

Gene-level GI scores were rearranged into a square symmetric matrix by ordering genes in rows and columns so that matrix positions i,j and j,i encode the same score for one gene pair. Scores for gene pairs that did not survive all filters, and were thus missing from the final dataset (62), were imputed using the 10-nearest neighbors through the impute package^157^ in R. The resulting matrices were clustered using average linkage hierarchical clustering with uncentered Pearson correlation using the hclust function in R. To produce the heatmap enriched for supported, high-confidence γGIs shown in Figure 2E, genes were subset according to the following procedure: first, we defined gene-level γGI scores as “supported” if at least 4 model-specific, sgRNA-level γGI were averaged to generate the score. Scores with this classification increased confidence by ensuring that more than a single pair of sgRNAs (one or both orientations; *i.e.*, A/B and B/A) contributed to the measurement (Figure S3E; see “Calculating gene-level GI scores”). Second, we defined “high-confidence” γGI scores as those with local FDR ≤ 0.01. Third, we computed the number of “supported high-confidence interactions” for each gene in our dual-sgRNA library and selected genes with at least the median number (determined among genes with at least one supported high-confidence interaction). Although heatmaps of τGI scores were not displayed in the manuscript, we followed the same procedure for these data and included annotations of genes enriched for supported, high-confidence interactions in Table S4. Heatmaps throughout the paper were produced using the R package ComplexHeatmap.^158^

To annotate functional modules defined by our interaction map, we used weighted correlation network analysis, implemented via R package WGCNA.^159^ This approach combines a soft thresholding of noisy scores by raising the distance matrix to higher powers with dynamic branch cutting^160^ applied to the hierarchical tree that results from clustering the thresholded distances. Soft thresholding powers were chosen to produce a set of “medium- and high-stringency clusters” by identifying those which best approximated a scale free network topology, one where a few highly connected genes act as hubs linking less-critical genes to the system. We used this methodology to identify functional modules among all genes as well as a subset enriched for supported, high-confidence interactions using both γGI and τGI scores. For medium-stringency clusters, we used soft thresholding powers of 3 and 4 for γGI scores (all genes and enriched subset, respectively) and 4 and 6 for τGI scores; for high-stringency clusters, 7 and 6 were used for γGI scores (all genes and enriched subset, respectively) and 9 and 15 were used for τGI scores (Table S4).

#### Considerations to guide data exploration

Our data offer insights into the vast array of mechanisms underlying genome stability and response to PARP inhibition. As such, we anticipate that these data will be mined for hypothesis generation and validation purposes. To support these efforts, we examined features of the dataset with the potential to confound interpretation of individual gene-level interactions. We provide discussion on two of these features below and one above (see “Calculating gene-level GI scores”). Any information relevant to interpretation is also annotated alongside those interactions (21,543) in Table S3.

#### Replicate variability

During our GI screen, the growth rate of one of our reference replicates diverged from the growth rate of the other temporarily, likely due to cell-handling stress (Figure S1E). Correspondingly, for some sgRNAs and sgRNA pairs in the reference condition, we observed a difference between replicate phenotypes (Figure S3F). More specifically, phenotypes of 19 individual sgRNAs (see “Calculating construct and sgRNA phenotypes”) were significantly different between replicates, as determined by Rosner’s generalized extreme Studentized deviate test (Rosner’s Test, implemented in R package EnvStats;^161^ Figure S3F). As we had sufficient data points that were approximately normally distributed, Rosner’s Test was preferred for its ability to identify multiple outliers simultaneously. Because genes targeted by these sgRNAs were enriched for homology directed repair (FDR = 1.82e-02, Reactome pathways; Table S2), with a single *BRCA2*-targeting sgRNA the strongest outlier, we examined γGI scores associated with these genes carefully. Indicating that gene-level scores were robust to phenotype variability across replicates, none deviated from the expected relationship between γGI score replicate correlation and the number of significant interactions per gene (Figure S3G).

#### Poorly performing object sgRNAs

When evaluating the robustness of gene-level γGI scores (Figure S3G), we identified 13 genes with an abnormally high number of significant interactions relative to γGI score replicate correlation (Rosner’s Test). One such gene, *NUP62*, had 202 significant interactions, 50 more than any other gene. To understand what was driving this effect, we examined this gene. First, as one *NUP62*-targeting sgRNA was removed from the dataset during initial filtering, the maximum number of supporting sgRNA-level γGI for each gene paired with *NUP62* was reduced to four, thus decreasing the reliability of resulting scores (Figure S3E; see “Calculating construct and sgRNA phenotypes” and “Calculating gene-level GI scores”). Next, we defined the model generated by querying the surviving *NUP62*-targeting sgRNA as the “*NUP62* query model”, and models generated with the partner sgRNA for each sgRNA pair with *NUP62* as “*NUP62* object models”, then contrasted the scores called by these models.

Indicative of some technical artifact, the *NUP62* query model did not show a convincing relationship between individual sgRNA and combinatorial phenotypes (Figure S3H). Moreover, phenotypes associated with non-targeting pairs had a wide distribution, compressing the resulting normalized interaction scores (see “Calculating model-specific, sgRNA-level GI scores.”). Indeed, if gene-level γGI scores were calculated using just this model, only 22 would be called significant. The query model was therefore not driving the aberrantly high number of significant scores. By contrast, although generally correlated with query model scores, sgRNA-level γGI scores called for *NUP62* interactions from object models had high variance with stronger scores (Figure S3I). We attribute the abnormally high number of significant gene-level γGI scores to these models and, further, attribute poor performance of this sgRNA as an object to the relatively static phenotypes observed across pairs (Figure S3H). Since we interpret these phenotypes to be artifactual, most γGIs with *NUP62* are also expected to be artifacts. Given this observation, we identified other sgRNAs with similar features and annotated any gene-level γGI and τGI scores that may be affected (15,552) in Table S3.

Notably, read counts for the constructs involving the 35 sgRNAs identified as poor objects across both the γ and τ phenotypes were found to be less than half that of the mean representation at T0 (117 vs 258).

#### Additional resources

Interaction screen data can be interactively explored through our accompanying web application at https://parpi.princeton.edu/map.

## SUPPLEMENTARY FIGURES AND LEGENDS

**Figure S1.**
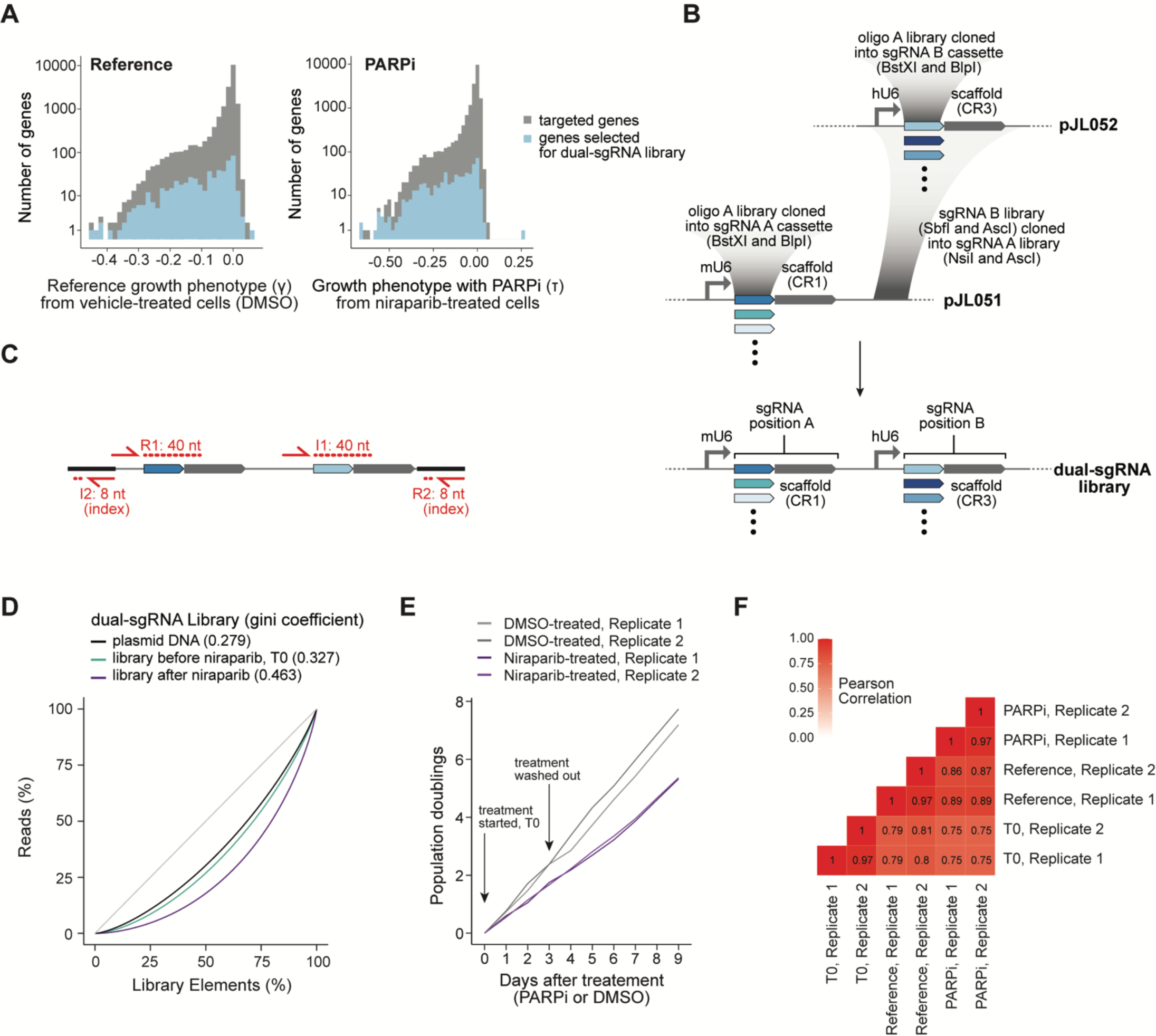
Designing, building, and screening a dual-sgRNA CRISPRi library, related to Figure 1. (A) Distribution of GI library gene growth phenotypes from single-perturbation genome-scale screen. (B) Dual-sgRNA CRISPRi library cloning strategy (Methods). (C) Dual-sgRNA library sequencing strategy (Methods). (D) Lorenz curves of the distribution of dual-sgRNA library elements at different experimental points. The gray line indicates a theoretical library in which every element appears with the same frequency. Gini inequality coefficients are shown in parentheses, with smaller coefficients indicating a more even distribution of library elements. (E) Population doublings during interaction screen for vehicle-(gray) and niraparib-treated (purple) replicates. (F) Pearson correlations of counts at start of interaction screen (T0).

**Figure S2.**
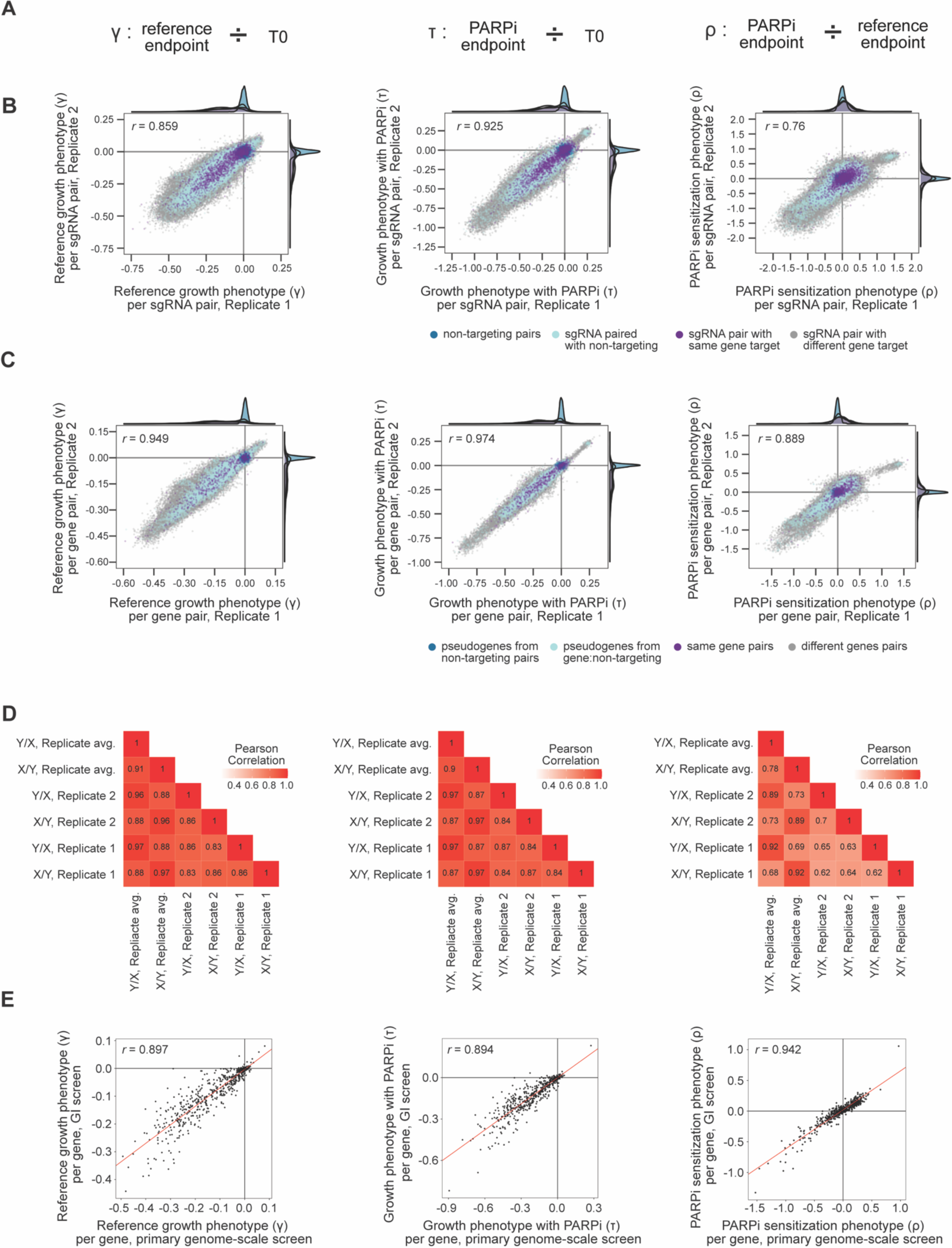
Phenotype reproducibility from single- and double-perturbation across conditions, related to Figure 2. (A) Conceptual definition of gamma (γ, left), tau (τ, middle), and rho (ρ, right) phenotypes (Methods). (B) γ (left), τ (middle), and ρ (right) sgRNA pair phenotypes across independent replicates. Pearson correlations are indicated. (C) Gene pair phenotypes across independent replicates. As in B. (D) Pearson correlations across orientations (X/Y vs Y/X) for independent replicates and the replicate average construct phenotypes. Only sgRNA pairs which had both orientations survive low frequency filters (Methods) were used in this analysis. As in B. (E) Cross-screen gene phenotype comparison. As in B.

**Figure S3.**
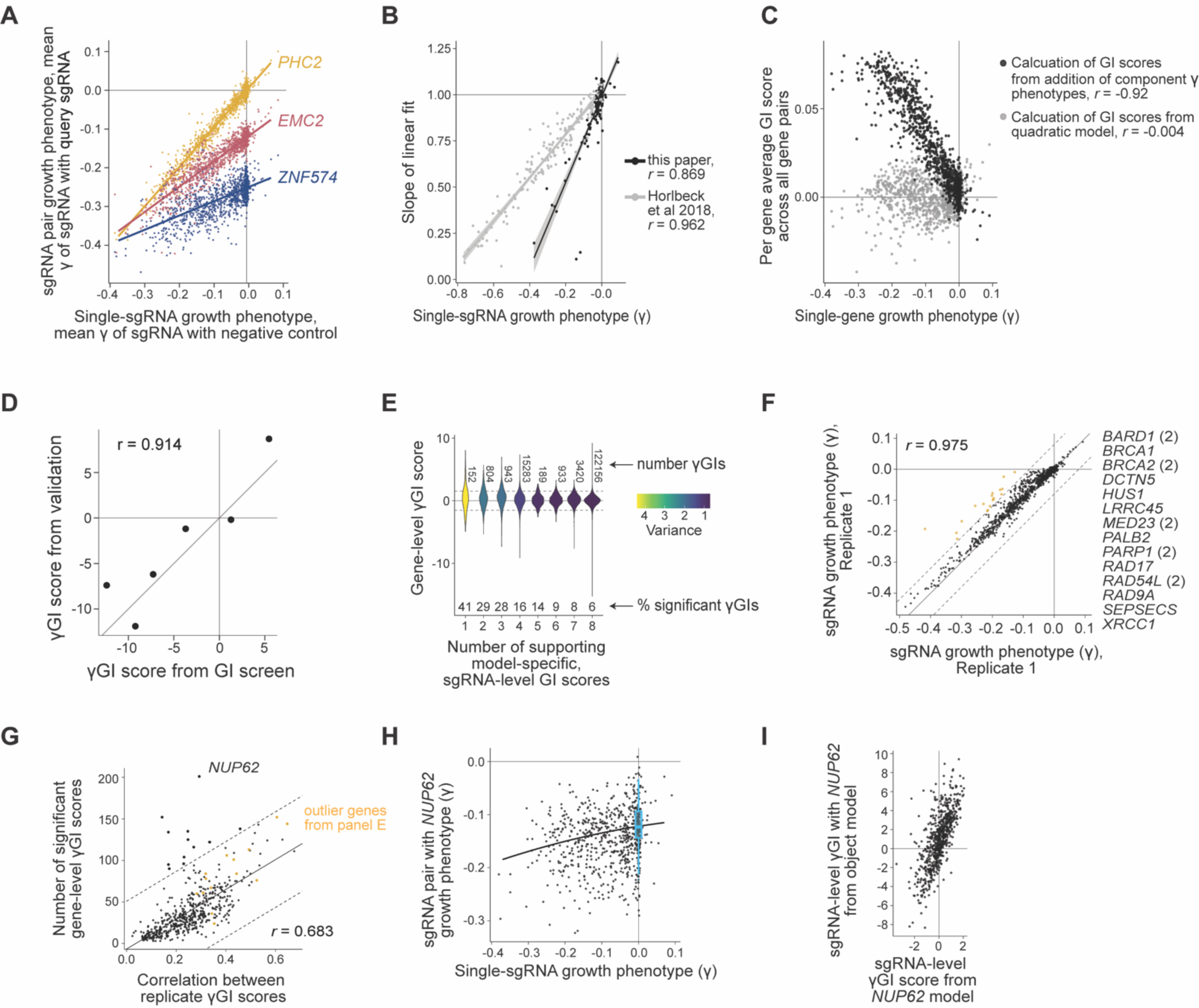
Considerations to guide data exploration, related to Figure 2. (A) Models for sgRNAs targeting *PHC2* (yellow), *EMC2* (red), and *ZNF574* (blue). In each case, a linear model (lines) outperformed a quadratic model. (B) Relationship between the degree of correction (model slope) for a query sgRNA and its γ growth phenotype. Data shown for all sgRNAs for which a linear fit worked well, in our screen (black) and a separate published interaction screen (gray).^48^ Pearson correlations are shown. (C) Average unnormalized γGI score for each gene across all pairs involving that gene against the associated single-gene γ phenotypes. Expected combinatorial phenotypes were calculated as either the sum of contributing phenotypes (black) or via a quadratic fit (gray; Methods). Pearson correlations are shown. (D) γGI score comparison, GI screen against low-throughput validation experiments (Methods). Pearson correlation is shown. (E) Distribution of γGI scores by number of supporting interactions (Methods). Dotted lines indicate significance thresholds corresponding to FDR ≤ 0.05. (F) Individual sgRNA γ phenotypes across independent replicates. sgRNAs in yellow were identified by Rosner’s test as having significantly different phenotypes across replicates (threshold indicated by dotted lines). Genes targeted by identified sgRNAs are listed; (2) indicates that both sgRNAs targeting that gene were identified. (G) Number of significant γGI called per gene against the cross-replicate correlation of its γGI scores. Dotted lines indicate boundaries of outliers according to Rosner’s test, and genes from the analysis in F are indicated in yellow. (H) *NUP62* guide model. Model fit is shown as the black line and distribution of non-targeting guides is shown in blue (boxplot). (I) Comparison of *NUP62* γGI scores called from the *NUP62* query and object sgRNA models.

**Figure S4.**
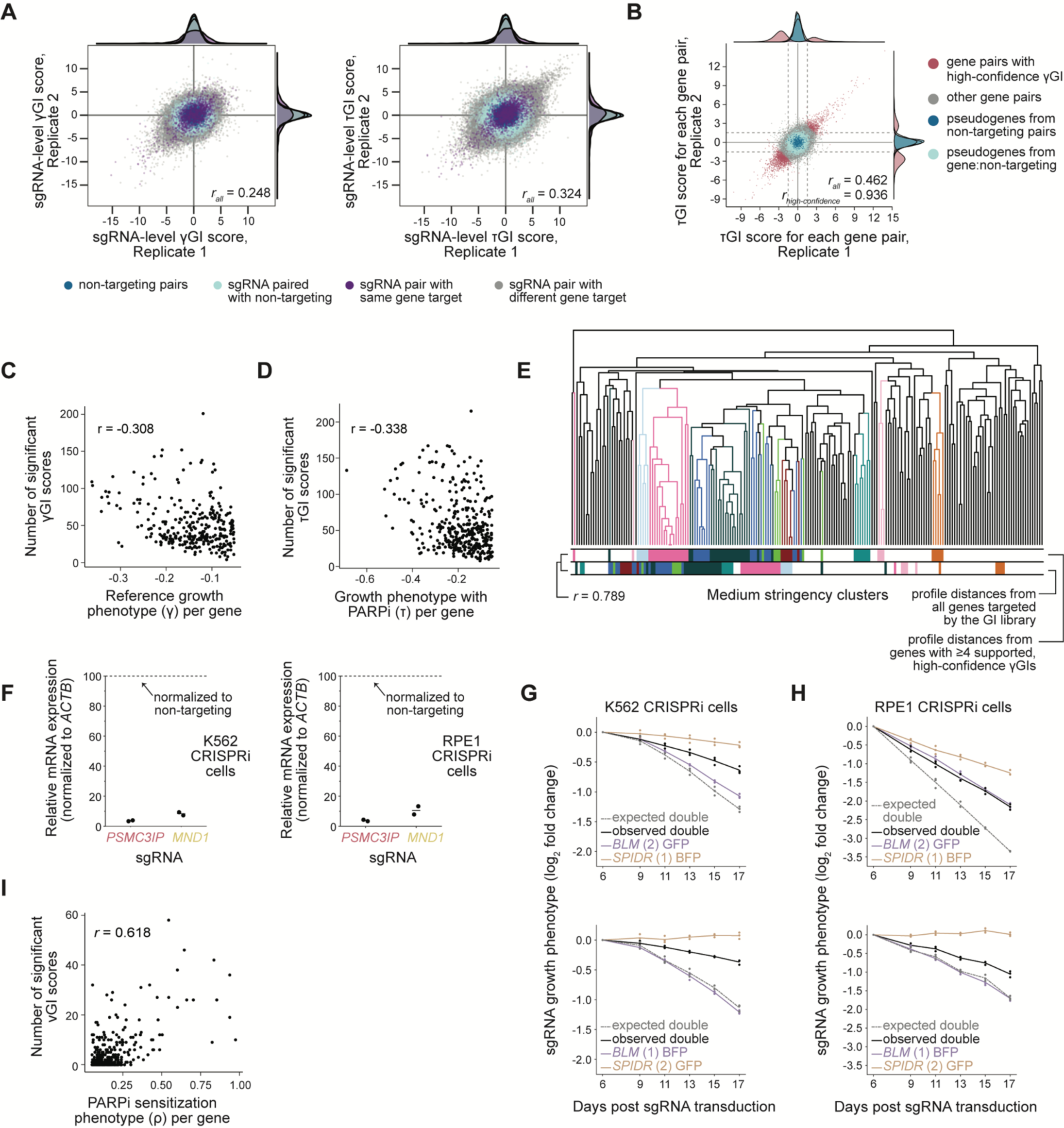
Analysis and validation of genetic interaction scores, related to Figures 3 and 4. (A) sgRNA pair γGI (left) and τGI (right) scores across independent replicates. Scores are the average of up to two model-specific, sgRNA-level GI scores (Methods). Pearson correlations are indicated. (B) Gene-level τGI score measurements for independent replicates. Dotted lines indicate significance bounds corresponding to FDR ≤ 0.05. Pearson correlations are given for all interactions and for high-confidence interactions. (C-D) Relationship between the number of significant γGI (C) and τGI (D) identified for a gene and its individual γ or τ phenotype. Only genes with growth phenotype γ, τ ≤ -0.05 were included, and Pearson correlations are listed. (E) Hierarchical clustering comparison using either distances calculated from only genes in the high-confidence enriched submap presented in Figure 2E or from distances calculated using all genes in the GI library. (F) qPCR confirms knockdown of *PSMC3IP* and *MND1* by both sgRNAs used for validation experiments in K562 CRISPRi (left) and RPE1 CRISPRi (right) cell lines. ACTB served as the reference gene. Cells transduced with non-targeting sgRNAs served as reference samples. ΔΔCt value was calculated and converted to percentage of expression level. (G-H) Validation of buffering interaction between *BLM* and *SPIDR* with two sets of sgRNAs in K562 CRISPRi (G) and RPE1 CRISPRi (H) cell lines. As in Figure 3D. (I) Relationship between the number of νGI hits identified for a gene and the magnitude of its ⍴ phenotype. Only genes with PARPi sensitization phenotype |⍴| ≥ 0.05 were included; *PARP1* was excluded as an outlier. Pearson correlation is listed.

**Figure S5.**
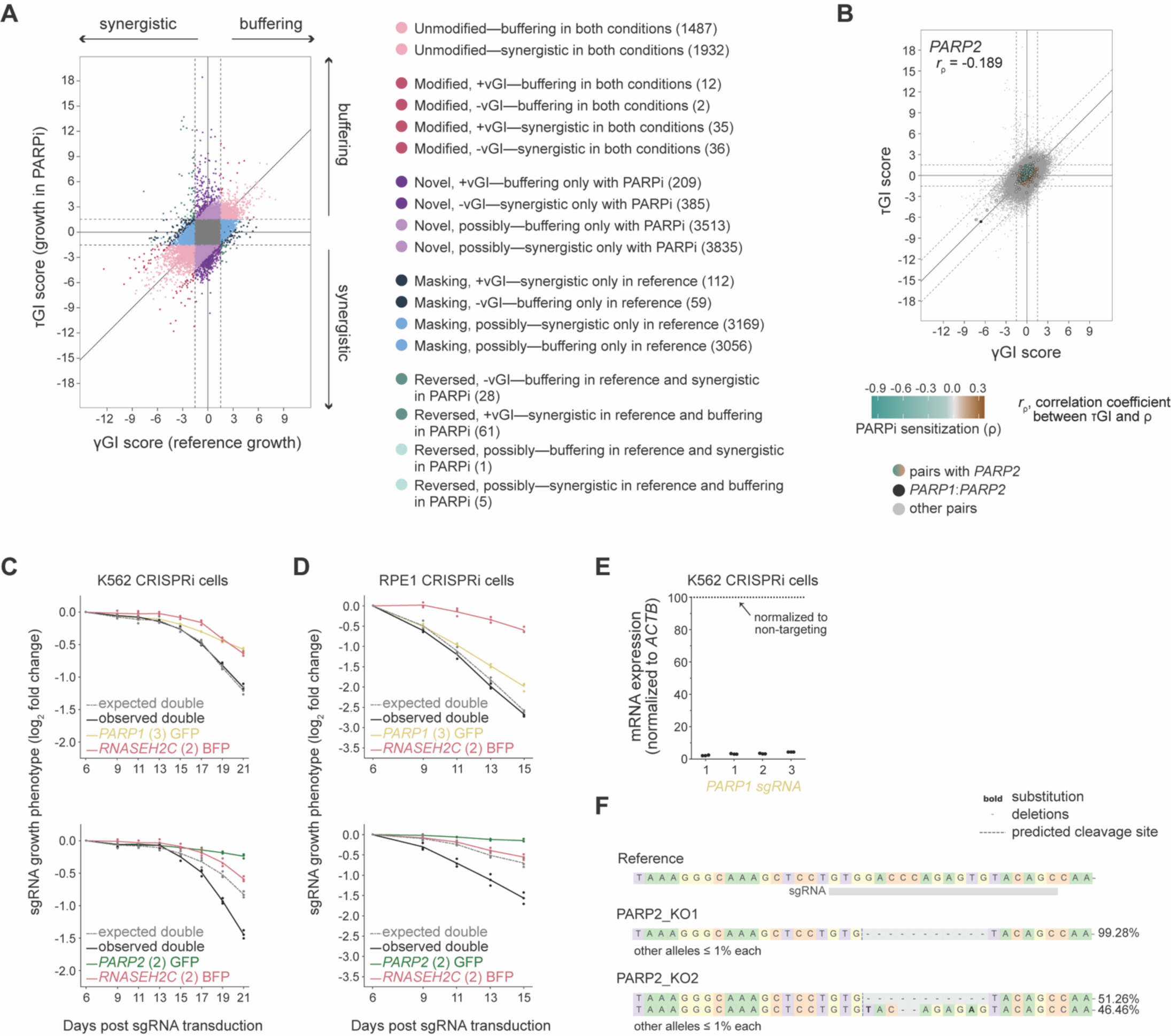
Expanded interaction topology and additional validation, related to Figures 4 and 5. (A) Expanded interaction category definitions (Methods; Tables S3 and S5). Dotted lines define significance thresholds corresponding to FDR ≤ 0.05. (N) after each category definition indicates the number of observed interactions in that category. (B) *PARP2* interaction topology. As in Figure 4C. (C-D) Additional validation experiments for the *PARP1*:*RNASEH2C* (top) and *PARP2*:*RNASEH2C* (bottom) interactions in K562 CRISPRi (C) and RPE1 CRISPRi (D) cell lines. As in Figure 3D. (E) qPCR confirms knockdown of *PARP1* sgRNAs expressed from either BFP or GFP tagged single-perturbation vector in K562 CRISPRi cells. ACTB served as the reference gene. Cells transduced with non-targeting sgRNAs served as reference samples. ΔΔCt value was calculated and converted to percentage of expression level. (F) Alleles from both *PARP2* knockout RPE1 CRISPRi clones at the targeted *PARP2* locus determined by sequencing. Reference sequence was from the GRCh38 assembly of the human genome.

**Figure S6.**
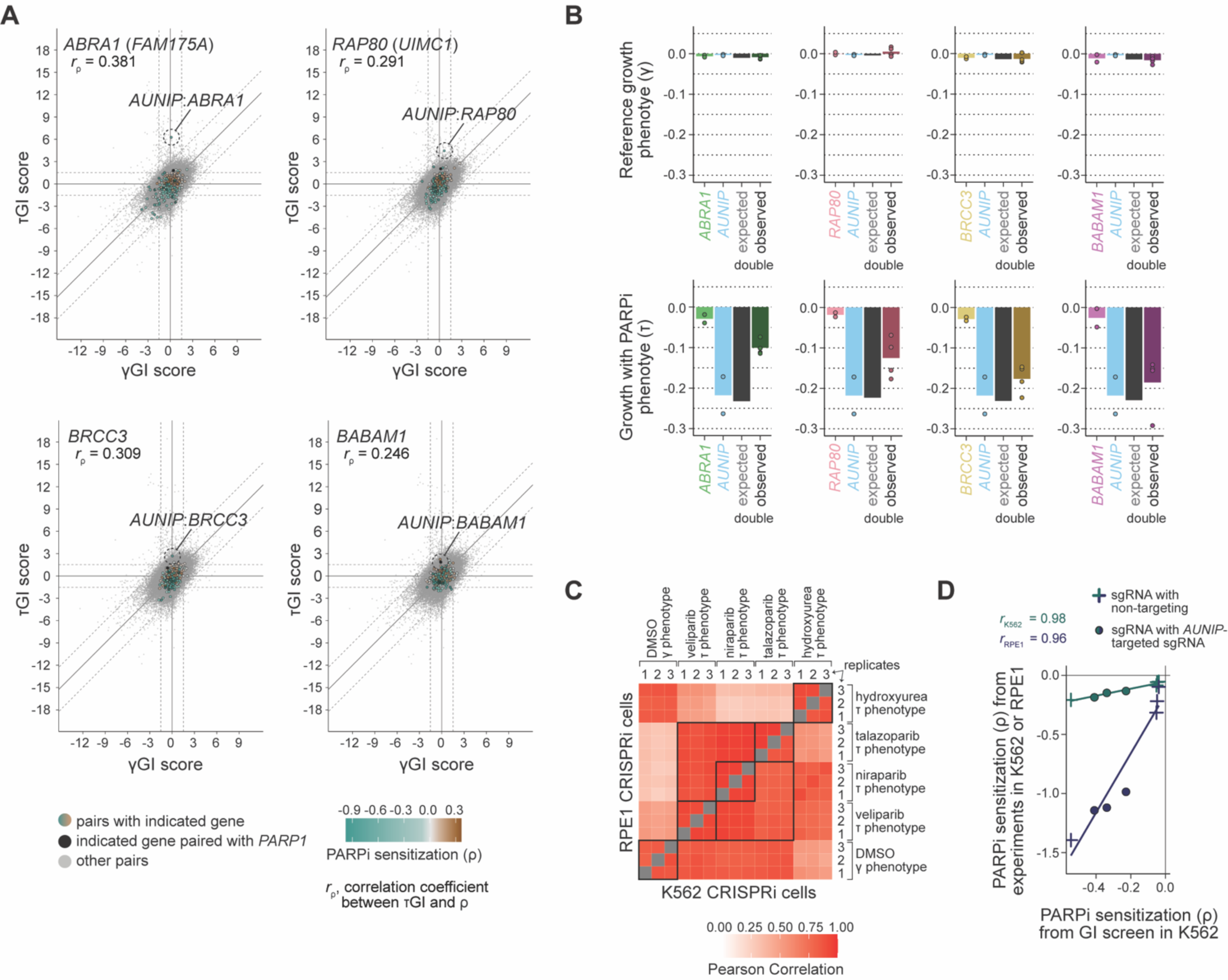
Analysis of interactions between *AUNIP* and BRCA1-A complex genes, related to Figure 6. (A) Interaction topologies for *ABRA1* or *FAM175A* (top left), *RAP80* or *UIMC1* (top right), *BRCC3* (bottom left), and *BABAM1* (bottom right). As in Figure 4C. (B) Observed single and combinatorial growth phenotypes from interaction screen in reference (top) and PARPi-treated conditions (bottom) for *AUNIP* (blue) with *BRCC3* (yellow), *RAP80* or *UIMC1* (red), and *ABRA1* or *FAM175A* (green). As in Figure 3C. (C) Growth phenotype replicate correlations with and without presence of drug in *AUNIP* target screen. The upper left triangle contains correlations measured in RPE1 CRISPRi cells, with K562 CRISPRi cell phenotype correlations in the bottom right. Black squares section phenotypes and drugs. (D) Comparison of niraparib ρ phenotypes observed in the interaction screen and in the *AUNIP* target screen by cell line (K562 CRISPRi, teal; RPE1 CRISPRi, navy). Colored lines indicate a linear fit for each cell line. Pearson correlations are listed.

## SUPPLEMENTARY TABLE TITLES AND LEGENDS

**Table S1. Single-perturbation genome-scale screen results, related to Figure 1**. Per sgRNA counts in each replicate at beginning of screen (T0) and at each endpoint (DMSO-treated and PARPi-treated). γ, τ, and ρ sgRNA phenotypes, along with gene-level p-values and q-values determined by MAGeCK are given. Additionally, annotation columns indicate whether an sgRNA was selected for inclusion in the GI library (“GI.Screen”), and if so, whether the guide survived all filters and generated gamma (“GI.Gamma”) or tau (“GI.Tau”) GI scores.

**Table S2. Results from all database enrichment searches performed, related to Figures 1-3 and 6**. Genes used for each search are listed (“Query genes”).

**Table S3. Genetic interaction scores, assigned interaction categories, and sgRNA-level considerations, related to Figures 2-6**. Gene-level γ, τ, and ν GI scores (“Gamma.GI”, “Tau.GI”, “Nu.GI”) with number of supporting model-specific, sgRNA-level interaction scores (“Nsupp”) are listed alongside significance values. Interaction categories and the number of contributing scores with an sgRNA identified as having either replicate variability in the gamma setting (“Gamma.RepVar”) or as a poor object sgRNA in either setting (“Gamma.PoorObject”, “Tau.PoorObject”) are given.

**Table S4. Gene clusters and phenotypes, related to Figures 2-6**. Gene-level γ, τ, and ρ phenotypes are provided beside assigned clusters identified using either all genes (“All”) or a subset enriched for high-confidence interactions (“Sel”, “Gamma.Selected”, “Tau.Selected”) under either a medium (“Medium”) or high (“High”) stringency soft thresholding (Methods).

**Table S5. Interaction category enrichments by gene, related to Figure 4**. P-value determined by Fisher’s exact test indicating whether each gene is enriched for interactions of the indicated types (Figure S5A). Pearson correlation between ρ phenotypes and τGI scores as in Figure 4C are provided (“Pearson.r”).

**Table S6. *AUNIP* and BRCA1-A target interaction screen results, related to Figure 6**. Per replicate and per cell line counts and phenotypes for each library construct are provided. Drug treatments used are veliparib (“VELIP”), niraparib (“NIRAP”), talazoparib (“TALAZOP”), and hydroxyurea (“HU”).

**Table S7. Oligonucleotide primer sequences.** Collected identifiers and sequences for primers used in this study.

